# Feedforward ribosome control mitigates gene activation burden

**DOI:** 10.1101/2021.02.11.430724

**Authors:** Carlos Barajas, Hsin-Ho Huang, Jesse Gibson, Luis Sandoval, Domitilla Del Vecchio

## Abstract

Heterologous gene activation causes non-physiological burden on cellular resources that cells are unable to adjust to. Here, we introduce a feedforward controller that increases ribosome level upon activation of a gene of interest (GOI) to compensate for such a burden. The controller achieves this by activating a modified SpoT enzyme with sole hydrolysis activity, which lowers ppGpp level and thus de-represses ribosomes. Without the controller, activation of the GOI decreased growth rate by more than 50%. With the controller, we could activate the GOI to the same level without a growth rate decrease. A cell strain armed with the controller in co-culture enabled persistent population-level activation of a GOI, which could not be achieved by a strain devoid of the controller. The feedforward controller is a tunable, modular, and portable tool that for the first time allows dynamic gene activation without growth rate defects for bacterial synthetic biology applications.

## Introduction

In bacterial synthetic genetic circuits, genes work in orchestration to accomplish a variety of functions, from monitoring stress level and releasing drugs in the gut [1, 2, 3, 4], to sensing environmental pollutants in soil or water [5, 6, 7, 8, 9, 10]. In these circuits, genes become dynamically activated and repressed, depending on the environment and on the circuit’s state. When a gene of interest (GOI) is activated, cellular resources that the cell would otherwise devote to growth are used by the GOI’s expression. This burden on cellular resources decreases growth rate and leads to physiological changes with poorly predictable outcomes that generally hinder the intended performance of the engineered cell [11, 12, 13, 14, 15, 16, 17]. Decreased growth rate upon a GOI activation has especially severe consequences when engineered bacteria are in co-culture with other strains. In fact, co-existence of multiple strains in co-culture is contingent on tightly matching growth rates, wherein small growth rate differences between the strains typically lead to extinction of the slower growing strain [18, 19, 20]. Therefore, growth rate reduction of an engineered bacterial strain upon a GOI activation, by leading to this strain’s extinction in co-culture, also leads to loss of the population-level expression of the GOI.

To mitigate the consequences of gene expression burden, researchers have devised methods that make synthetic gene expression robust to changes in the availability of cellular resources [21, 22, 23, 24, 25]. Complementary approaches have also used orthogonal ribosomes for heterologous expression through synthetic rRNAs [26]. However, none of these tools can control growth rate. The problem of controlling growth rate has been addressed by a feedback controller that senses gene expression burden and reduces the GOI’s expression to low levels such that growth rate is not affected [27]. While this approach is ideal to maximize protein yield in batch-production, it is not suitable when the GOI needs to be dynamically activated to a specific and possibly high level as in biosensors and genetic logic gates [28, 29].

Here, we introduce a feedforward controller that allows the activation of a GOI to any desired level while keeping growth rate constant. The controller co-expresses SpoTH, a modified version of SpoT with only hydrolysis activity, with the GOI. When the GOI is activated, SpoTH is also expressed and catalyzes the hydrolysis of ppGpp, thereby de-repressing ribosomal rRNA and increasing ribosome level [30, 31, 32]. To achieve sufficient de-repression of the rRNA in any bacterial strain, we induce the expression of RelA+, a variant of RelA protein that exhibits constitutive ppGpp synthesis activity [33, 34]. RelA+ expression results in elevated levels of ppGpp, which are then hydrolyzed by SpoTH. When the SpoTH ribosome binding site (RBS) is properly tuned, SpoTH activation increases ribosome levels to a point that exactly compensates for the growth rate burden caused by the GOI’s activation. The controller achieves this compensation in multiple strains, at different nominal growth rates, and also in co-culture.

## Results

### Growth rate actuation via SpoTH in strains with elevated ppGpp levels

During balanced exponential growth, ppGpp is the primary regulator of rRNA [30, 31, 32] and there is an inverse relationship between basal ppGpp levels and both rRNA transcription, the rate-limiting step in ribosome production, and growth rate [30, 35, 36, 37, 38]. The RelA/SpoT Homologue (RSH) proteins are responsible for catalyzing the synthesis and hydrolysis of ppGpp as shown in Fig. 1-a [39, 40, 41]. In particular, the SpoT enzyme is bifunctional with both synthesis and hydrolysis capabilities, with the latter dominating in exponential growth [42], while the RelA enzyme is monofunctional with sole synthesis activity. To actuate growth rate, we exogenously express a modified version of SpoT (Supplementary note 1) with only hydrolysis activity (SpoTH). Activation of SpoTH catalyzes the hydrolysis of ppGpp [43], which upregulates both ribosome production and growth rate (Fig. 1-a). The growth rate response as SpoTH level is varied depends on the level of basal ppGpp (Fig. 1-b and mathematical model in Supplementary notes 2 and 3).

**Figure 1:**
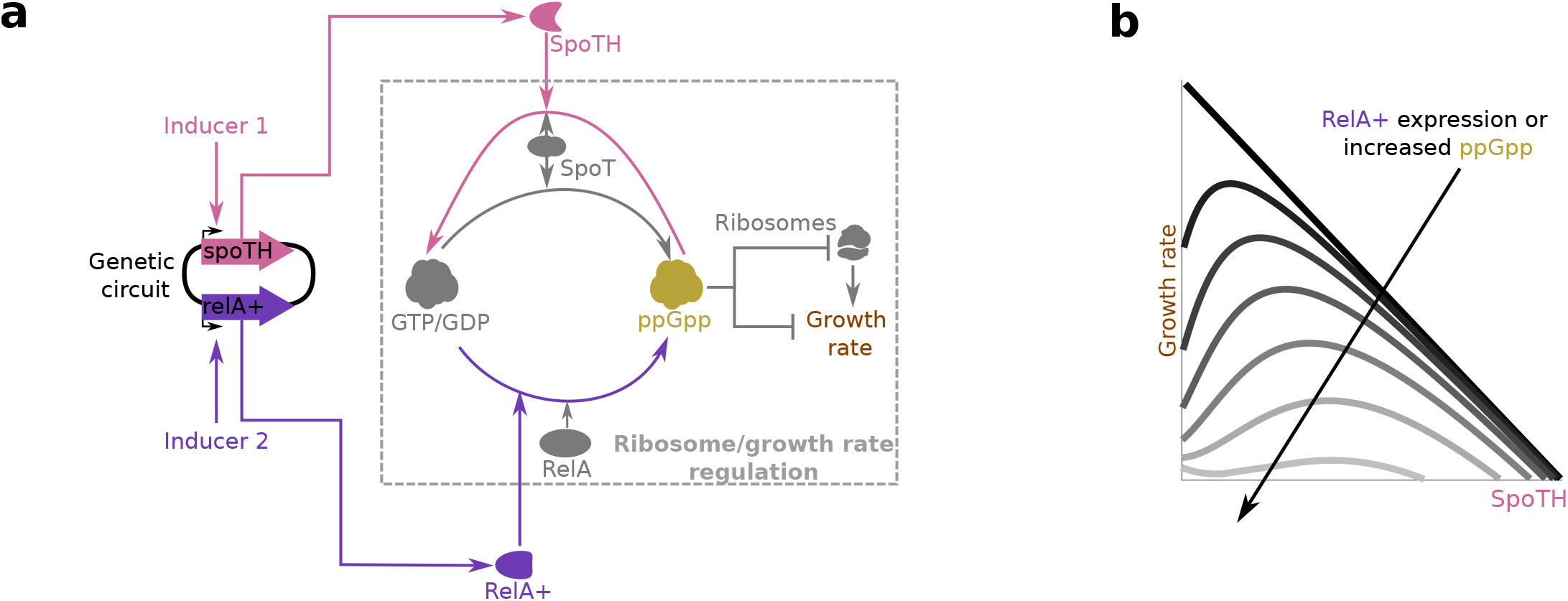
Growth rate actuation via SpoTH and RelA+. (**a**) Diagram of the circuit describing how SpoTH and RelA+ expression affect ribosomes and growth rate. The nucleotide ppGpp negatively regulates ribosome production and growth rate in *E. coli* during exponential growth. The synthesis of ppGpp is catalyzed from GTP/GDP by both RelA and SpoT, while the hydrolysis of ppGpp is catalyzed by SpoT only [39, 40, 41]. A modified version of SpoT with only hydrolysis activity (SpoTH) catalyzes the hydrolysis of ppGpp [43]. A modified version of RelA containing the N-terminal 455 residues of RelA (RelA+) catalyzes the synthesis of ppGpp [33, 34, 60]. Both SpoTH and RelA+ are expressed by a synthetic genetic circuit and are under the control of inducible promoters. (**b**) Growth rate as a function of SpoTH level for varying ppGpp concentration. SpoTH gene activation to low levels increases growth rate when the amount of ppGpp is sufficiently high. RelA+ expression increases ppGpp levels and thus decreases growth rate. See Supplementary notes 2 and 3 for a mathematical model.

We experimentally characterized the ability of SpoTH expression to actuate growth rate in three different strains carrying mutations in the SpoT gene, resulting in different basal levels of ppGpp. Specifically, we tested the CF944 (*spoT202* allele), CF945 (*spoT203* allele), and CF946 (*spoT204* allele) strains, where the basal ppGpp levels are lowest for CF944 and highest for CF946 [35, 44, 45, 31]. Alongside these strains, we also characterized the growth rate response to SpoTH expression in the wild-type MG1655 strain (WT). The genetic circuit used to express SpoTH in these strains is shown in Fig. 2-a,b. For CF945 and CF946, activation of the SpoTH gene increased growth rate by up to 80% and 60%, respectively (Fig. 2-c). For MG1655 and CF944, which have lower basal level of ppGpp, we were unable to positively actuate growth rate as the SpoTH gene was activated. Free ribosome concentration, indirectly measured through a GFP monitor (Supplementary note 4), increases when SpoTH is expressed. Specifically, GFP production rate increases for MG1655, CF944, CF945, and CF946, by 22%, 45%, 90%, and 65%, respectively, when SpoTH is expressed (SI Fig. 7). Strain CF945 provides the most growth rate and free ribosome level actuation and thus it is the strain we proceed with.

**Figure 2:**
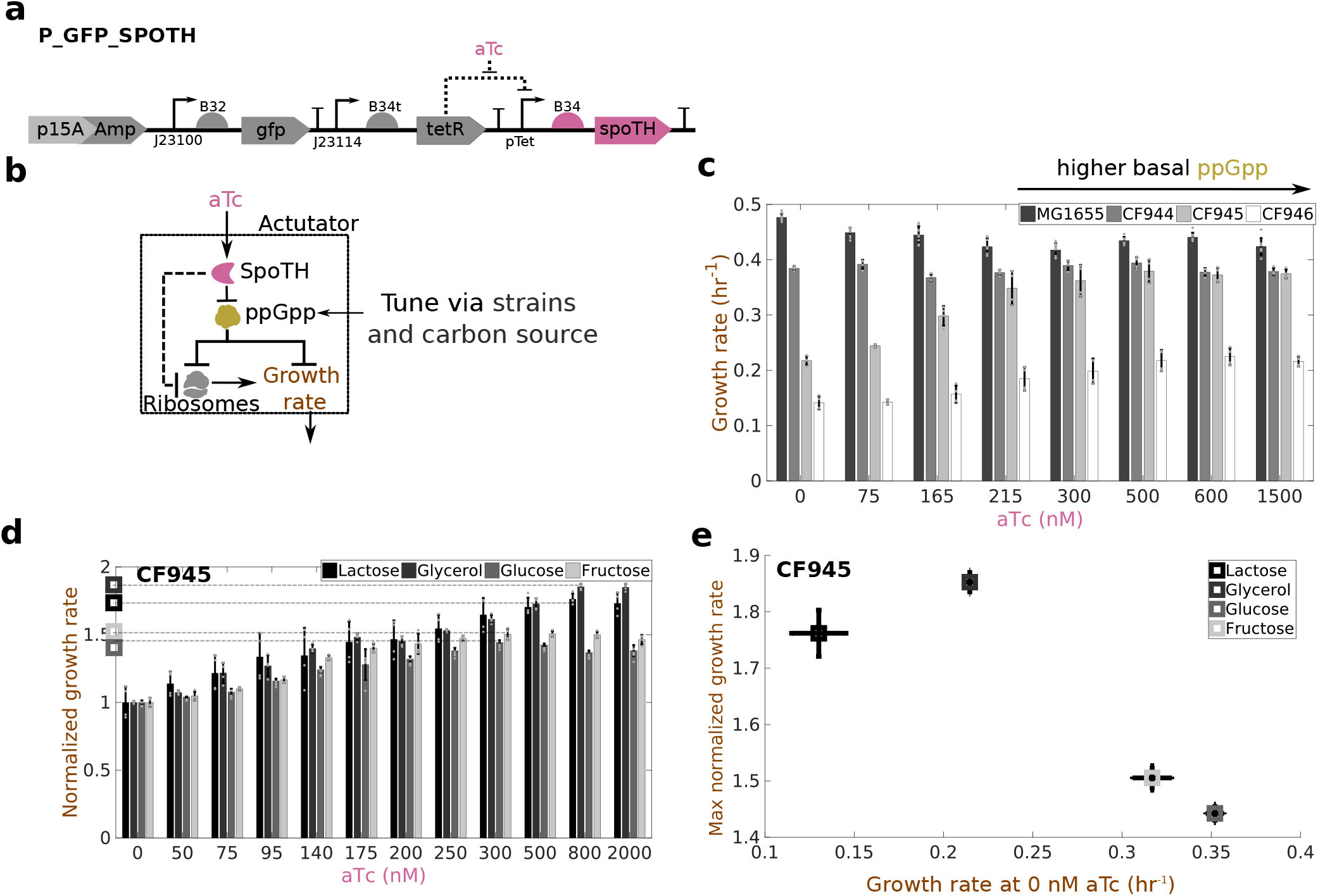
SpoTH gene activation increases growth rate. **(a)** P_GFP_SpoTH plasmid used to activate the SpoTH gene via the inducible pTet promoter. **(b)** Circuit describing how SpoTH induction via aTc affects ribosomes and growth rate. Addition of aTc increases SpoTH concentration, which lowers ppGpp concentration and consequently upregulates both free ribosome concentration and growth rate [35]. **(c)** The growth rate as SpoTH is increased in the wild-type MG1655, CF944, CF945, and CF946 strains [35] growing in glycerol as the sole carbon source. **(d)** Growth rate normalized by the growth rate at aTc = 0 nM, as SpoTH is expressed in CF945 growing in lactose, glycerol, fructose, or glucose as the sole carbon source. The maximum normalized growth rate for each carbon source is marked by open squares. **(e)** The maximum normalized growth rate versus the growth rate at aTc = 0 nM for each carbon source. Data are shown as the mean ± one standard deviation (N=4, two biological replicates each with two technical replicates). Individual experimental values are presented as gray dots. The complete experimental protocol is provided in the Materials and Methods section. Plasmid description, plasmid map, and essential DNA sequences are provided in SI section *Plasmid maps and DNA sequences*.

For a fixed strain, an additional method to tune basal ppGpp level is via the growth medium composition, specifically through the carbon source [32, 36, 46, 47, 48]. Consequently, we also tested four common carbon sources: glucose, glycerol, fructose, and lactose. The nominal growth rate without SpoTH expression was ∼ 0.35 hr^−1^, ∼ 0.32 hr^−1^, ∼ 0.2 hr^−1^, and ∼ 0.12 hr^−1^ and can be increased by up to ∼ 45%, ∼ 50%, ∼ 85%, and ∼ 75% by expressing SpoTH with glucose, fructose, glycerol, and lactose, respectively (Fig. 2-d,e). GFP production rate increases by ∼ 55%, ∼ 60%, ∼ 100%, and ∼ 150% when expressing SpoTH with glucose, fructose, glycerol, and lactose as the carbon sources, respectively (SI Fig. 7).

These data indicate that there is a tradeoff between nominal growth rate and the relative growth rate increase that can be achieved by SpoTH expression (Fig. 2-e). This tradeoff occurs because the extent to which growth rate can be increased is directly tied to the amount of ppGpp available to be hydrolyzed. That is, high basal ppGpp, yielding lower basal growth rate, allows for larger growth rate increase upon SpoTH expression (SI Fig. 19 and Supplementary note 3).

### Feedforward control of ribosomes in the CF945 strain

The feedforward ribosome controller co-expresses SpoTH with the red fluorescent protein (RFP) GOI (Fig. 3-a). We refer to this system as the closed loop (CL) system. The open loop (OL) system is a configuration where SpoTH is missing (Fig. 3-b). Addition of AHL activates the RFP gene, which sequesters ribosomes and negatively affects growth rate (upper branch in Fig. 3-c). In the CL system, however, addition of AHL also increases SpoTH expression (lower branch in Fig. 3-c), which increases ribosome level and growth rate, thereby compensating for the growth rate reduction caused by RFP gene activation. The mathematical model predicts that if the ribosome binding site (RBS) of SpoTH is appropriately tuned, then the availability of ribosomes increases exactly to match the demand by RFP gene activation (Fig. 3-d and Supplementary notes 2 and 3). Therefore, we designed four SpoTH RBS’s for the CL system with varying strengths (Supplementary note 4).

**Figure 3:**
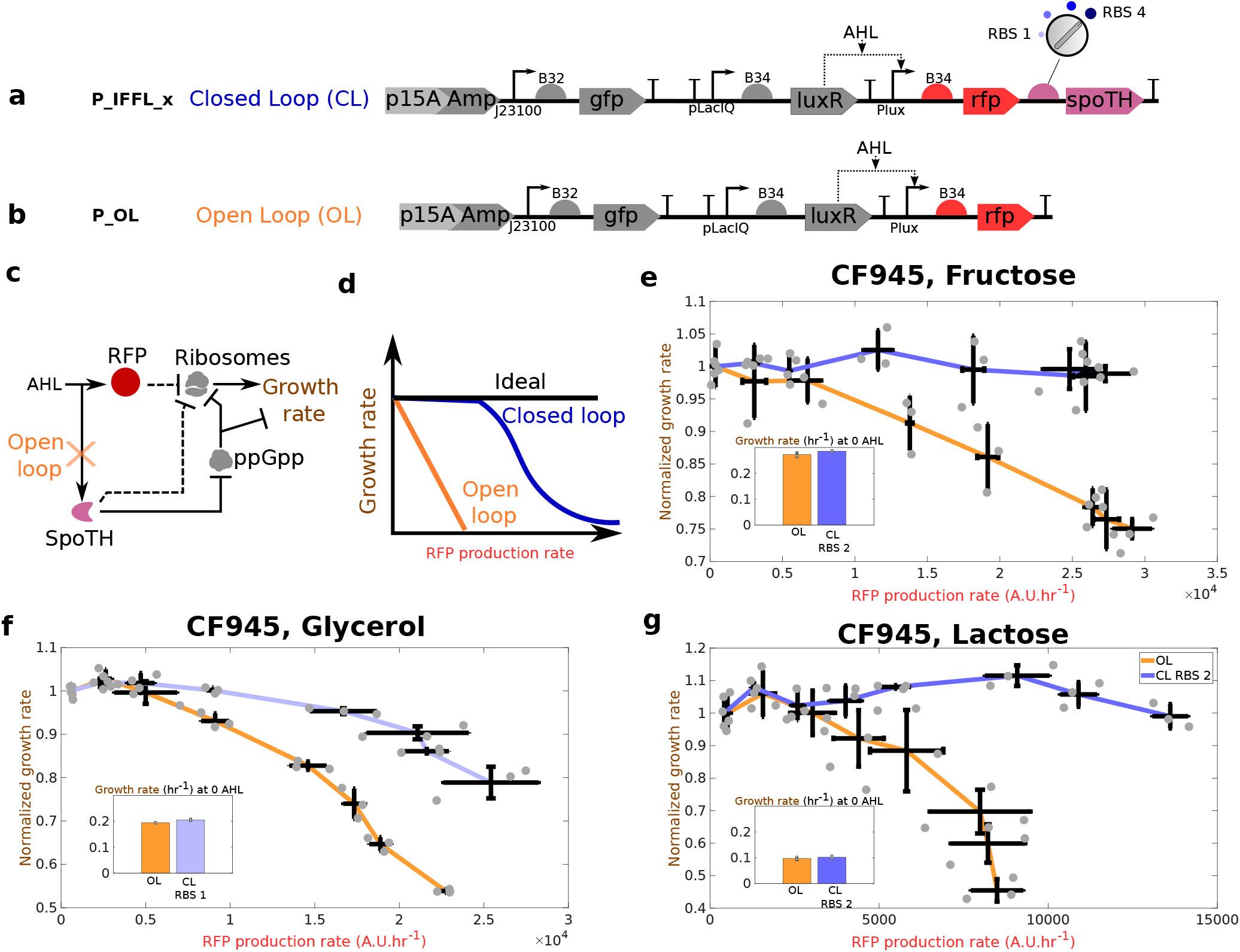
Feedforward ribosome controller compensates for burden caused by a GOI (RFP) activation at different nominal growth rates. **(a)** CL system’s genetic construct (P_IFFL___x) co-expresses RFP and SpoTH via the AHL inducible Plux promoter. The SpoTH RBS is used as a tuning parameter (Supplementary note 5). **(b)** OL system’s genetic construct (P_OL) expresses RFP using the AHL inducible Plux promoter. **(c)** Circuit diagram illustrating how AHL induction affects ribosomes and growth rate for the open loop (OL) or closed loop (CL) systems. In the OL system, SpoTH is not present, so there is only the upper path from AHL to ribosomes. In the CL system, AHL also activates SpoTH expression and hence upregulates ribosome concentration and growth rate. Dashed edges represent sequestration of free ribosomes by a protein’s expression. **(d)** Expected growth rate as the RFP gene is activated for the ideal case, the OL, and the CL systems. **(e-g)** Growth rate normalized by the growth rate at 0 nM AHL (nominal growth rate) versus the RFP production rate for the OL and CL systems, using fructose (e), glycerol (f), and lactose (g) as the carbon source. The inset shows the nominal growth rate with no AHL induction. Data for all the RBS’s of the CL system tested, AHL induction concentrations used, and GFP production data are shown in SI Fig. 8 and SI Fig. 9. Data are shown as the mean ± one standard deviation (N=3, three biological replicates). Individual experimental values are presented as gray dots. All experiments were performed in the CF945 strain. The complete experimental protocol is provided in the Materials and Methods section. Plasmid description, plasmid map, and essential DNA sequences are provided in SI section *Plasmid maps and DNA sequences*.

In fructose, the OL system growth rate drops by over 25% when activating the RFP gene, while for the CL system with RBS 2, the growth rate remains nearly constant when the RFP gene is activated to the same level (Fig. 3-e). In glycerol, the OL system growth rate drops by over 45% when activating the RFP gene, while for the CL system with RBS 1, the growth rate drops at most by 10% when we activate the RFP gene to the same level (Fig. 3-f). Finally, in lactose, the OL system growth rate drops by over 55% upon RFP gene activation, while for the CL system with RBS 2, the growth rate remains nearly constant for the same RFP gene activation (Fig. 3-g). The growth rate versus RFP production rate for other tested CL system’s RBS values is shown in SI Fig. 8. In lactose, cells have the lowest nominal growth rate and the largest actuation of GFP production rate when SpoTH is activated (SI Fig. 7). Consequently, for lactose, the CL system allows also GFP production rate to stay approximately constant when the RFP gene is activated, which is not possible in the OL system (SI Fig. 9).

We also considered a control genetic construct, where we replaced SpoTH with a nonfunctional heterologous protein CJB (cjBlue H197S [49]) (SI Fig. 10). This control construct allows to verify that the CL system outperforms the OL system due to the growth rate actuation by SpoTH expression and not because of the configuration change that the RFP mRNA undergoes when RFP is coexpressed with a second gene. This is confirmed since expressing RFP in this control circuit yields even lower growth rates than those of the OL system (SI Fig. 10). This is expected since CJB expression sequesters ribosomes adding to the burden of activating the RFP gene.

Taken together, these data indicate that the feedforward controller can be easily tuned across different nominal growth rates, which we achieved here by different carbon sources, to ensure no growth rate decrease upon the GOI’s activation.

### Feedforward control of ribosomes in common bacterial strain

To extend the feedforward controller to common bacteria, we introduced an inducible RelA+ gene expression cassette to elevate the ppGpp level in any strain of interest (Fig. 4-a,b). The *E. coli* RelA+ variant, containing the N-terminal 455 residues of wild type RelA protein, has constitutive ppGpp-synthesizing activity [33, 34]. The genetic construct used to express RelA+ and SpoTH is shown in Fig. 4-a,b. Increased levels of RelA+ in MG1655 (WT), TOP10, and NEB strains lead to increased level of ppGpp and hence to lower growth rate (Fig. 4-c). RelA+ expression also lowers GFP production rate consistent with down-regulation of free ribosome concentration (SI Fig. 11 and Supplementary note 4). For a level of RelA+ expression that halves the nominal strain growth rate, SpoTH gene activation upregulates growth rate close to the level with no RelA+ for all three strains (compare growth rate for maximal aTc in Fig. 4-d to that for no SAL in Fig. 4-c). Accordingly, the GFP production rate is also halved with RelA+ expression and upregulated back to nearly the value with no RelA+ expression through SpoTH gene activation (SI Fig. 11). We conclude that, with constitutive RelA+ expression, SpoTH gene activation allows to increase growth rate in common laboratory strains, thereby enabling transportability of the feedforward controller.

**Figure 4:**
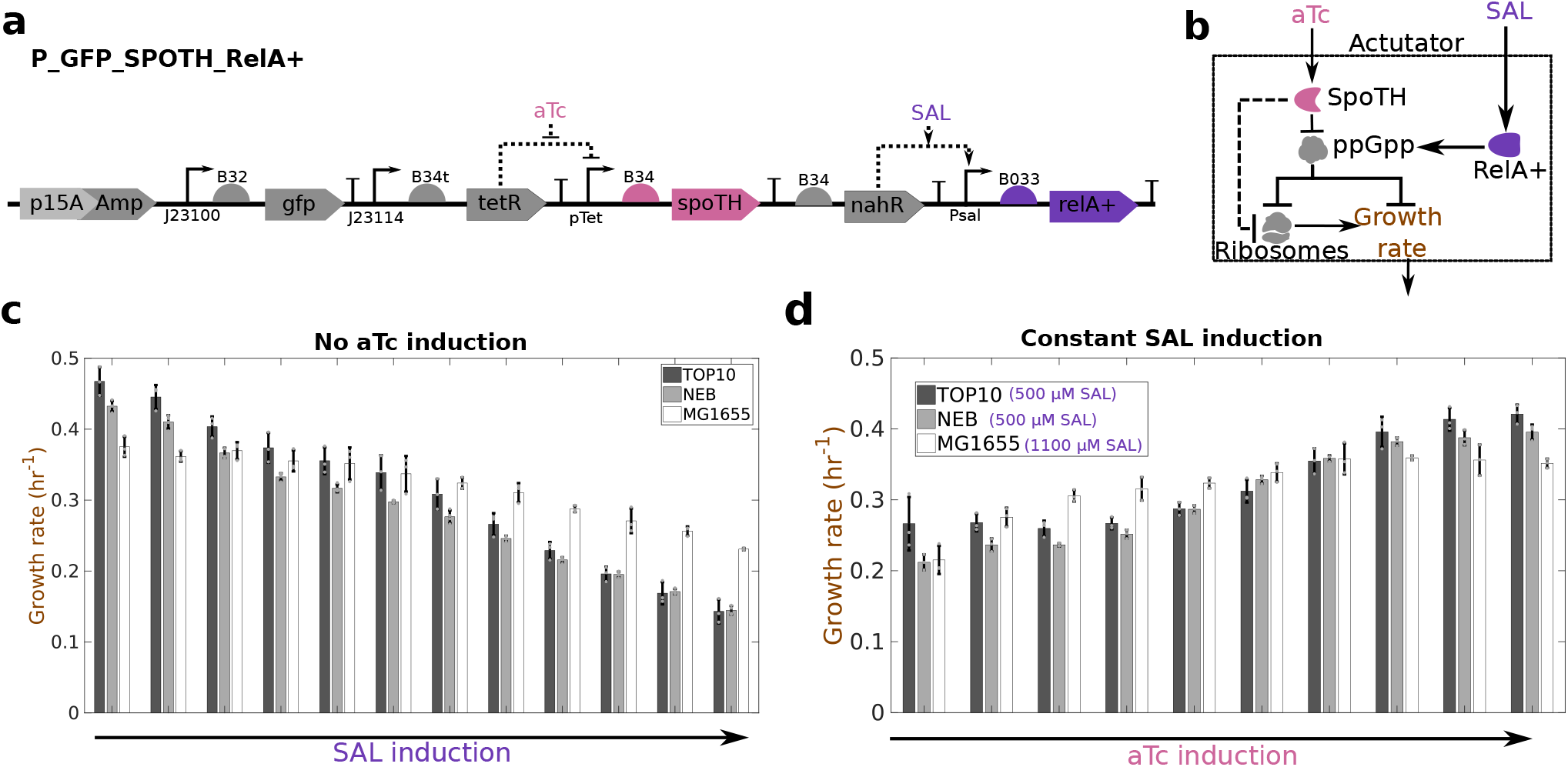
RelA+ expression allows to transport the ribosome controller to a desired bacterial strain. **(a)** P_GFP_SpoTH_RelA+ construct expresses SpoTH via the inducible pTet promoter and RelA+ via the inducible Psal promoter. Plasmid description, plasmid map, and essential DNA sequences are provided in SI section *Plasmid maps and DNA sequences*. **(b)** Circuit diagram depicting the effect of RelA+ induction and SpoTH induction on ribosomes and growth rate. Addition of SAL increases RelA+ concentration and thus upregulates ppGpp concentration [60]. Addition of aTc increases SpoTH concentration, which lowers ppGpp concentration and consequently upregulates both free ribosome concentration and growth rate [35]. **(c)** Growth rate versus RelA+ induction (SAL) in the TOP10, NEB, and wild-type MG1655, strains growing in glycerol as the sole carbon source. The SAL inductions are [0, 5, 10, 40, 75, 150, 250, 375, 750, 1000] *μ*M for the NEB and TOP10 strains and [0, 10, 20, 30, 50, 100, 175, 250, 375, 500, 750, 1000] *μ*M for MG1655 strain. **(d)** Growth rate versus SpoTH induction (aTc) for a fixed RelA+ induction in TOP10, NEB, and wild-type MG1655 strains growing in glycerol as the sole carbon source. The aTc inductions are [0, 20, 30, 40, 45, 50, 60, 70, 80, 90] nM for TOP10, [0, 20, 30, 40, 50, 60, 70, 80, 90, 100] nM for NEB, and [0, 40, 80, 120, 160, 200, 240, 280, 320, 360] nM for MG1655. Data are shown as the mean ± one standard deviation (N=3, three biological replicates). Individual experimental values are presented as gray dots. The complete experimental protocol is provided in the Materials and Methods section.

We next evaluated the ability of the feedforward controller to keep growth rate constant as the RFP gene is activated in a TOP10 strain (Fig. 5-a,b). To this end, we established three OL systems at different nominal growth rates by transforming the OL system circuit of Fig. 3-a in CF944, CF945, and CF946. We then evaluated three genetically identical CL systems all using RBS 2 (Fig. 5-a), each with nominal growth rate matching that of the corresponding OL system, which we obtained by adjusting the RelA+ expression level (insets of Fig. 5-c,d,e). This way, both OL and CL systems have matching growth rates before the RFP gene is activated.

**Figure 5:**
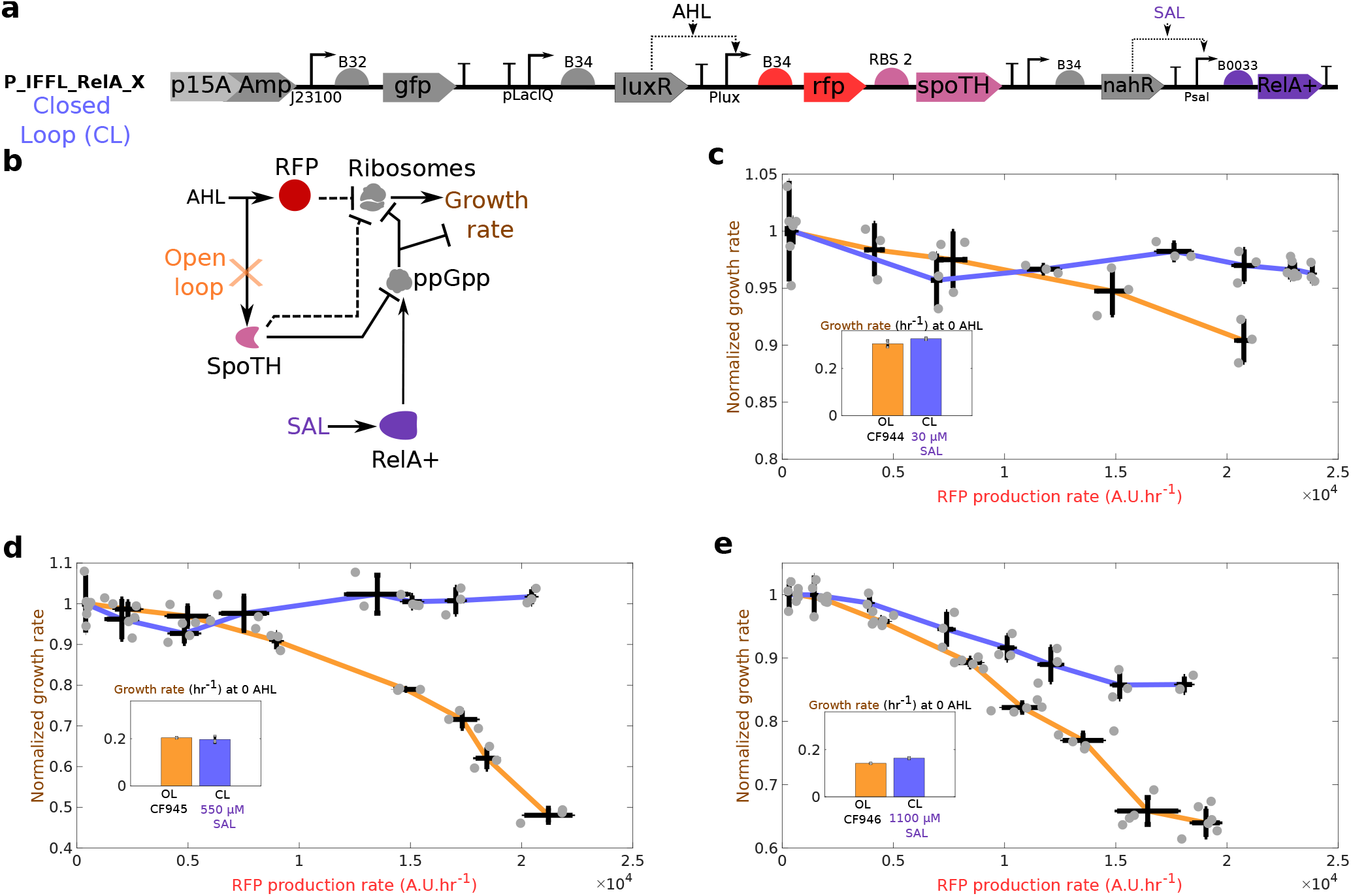
Feedforward ribosome controller compensates for burden caused by activation of the RFP gene in a common laboratory strain and across different nominal growth rates. **(a)** CL system’s genetic construct (P_IFFL_RelA_2) co-expresses RFP and SpoTH via the AHL inducible Plux promoter. The SpoTH RBS is fixed to RBS 2 (Supplementary note 5). RelA+ is expressed using the SAL inducible Psal promoter. **(b)** Circuit diagram depicting the effect of activating RFP (AHL input) on ribosomes and growth rate for the open loop (OL) or closed loop (CL) systems. In the OL system, SpoTH is not present, so there is only the upper path from AHL to growth rate. In the CL system, AHL also activates SpoTH production and hence upregulates ribosome concentration and growth rate. Dashed edges represent sequestration of free ribosomes by a protein’s expression. RelA+ activation via SAL sets the basal level of ppGpp and thus the nominal growth rate [60]. **(c-e)** Growth rate normalized by the growth rate at 0 nM AHL (nominal growth rate shown in the inset) versus the RFP production rate for the OL in CF944 (c), CF945, (d), and CF946 (e) and CL systems in TOP10. For the CL system, RelA+ expression is set to match the growth rate of the OL strain. The inset shows the nominal growth rate with no AHL induction. Data for all RBS’s of the CL system tested, AHL induction concentrations used, and GFP production data are shown in SI Fig. 12 and SI Fig. 13. Data are shown as the mean ± one standard deviation (N=3, three biological replicates). Individual experimental values are presented as gray dots. All experiments were performed with glycerol as the sole carbon source. The complete experimental protocol is provided in the Materials and Methods section. Plasmid description, plasmid map, and essential DNA sequences are provided in SI section *Plasmid maps and DNA sequences*.

When the RFP gene is activated, the growth rate of the OL system drops by 20%, 55%, and 40% in the CF944, CF945, and CF946 strains, respectively (Fig. 5-c,d,e). In contrast, the growth rates of the associated CL systems, only drop by 5%, 7%, and 15%, respectively, when the RFP gene is activated to the same level (Fig. 5-c,d,e). The RBS of the CL system in Fig. 5-e, can be further tuned to prevent a growth rate drop as the RFP gene is activated (Supplementary note 6). The growth rate versus RFP production rate for all CL RBS values tested is shown in SI Fig. 12. The GFP production rate versus the RFP production rate data corresponding to Fig. 5-c,d,e also demonstrates that for sufficiently low nominal growth rates, the SpoTH RBS can be tuned in the CL system to keep GFP production rate constant as the RFP gene is progressively activated (SI Fig. 13).

Taken together, these data show that RelA+ expression sets the nominal desired growth rate for the CL system in a strain of interest, and that the SpoTH co-activation with the GOI maintains this pre-set nominal growth rate as the GOI is activated.

### Feedforward controller for persistent GOI expression in co-culture

Engineered bacteria that dynamically express a GOI are often deployed in environments where other microbes are already present. Examples include engineered bacteria delivering biotheaputics in the gut microbiome or acting as biosensors for water contaminants [4, 7]. If the activation of the GOI leads to growth rate defects, then environmental faster growing organisms will overtake the population, leading to loss of the GOI population-level expression [18, 20]. This, in turn, hinders the sensing or drug delivery functionality of the engineered cell strain. Similarly, in engineered consortia, where multiple strains are programmed to each accomplish a different but complementary function, the different strains’ growth rates should remain sufficiently close to one another despite dynamic activation of genes [17, 50, 51]. Here, we tackle this problem by employing the feedforward controller to activate the GOI such that the strain’s growth rate does not change upon GOI activation.

Specifically, we compare the performance of the OL strain expressing inducible RFP (GOI) to that of the CL strain armed with the feedfroward controller, when co-cultured with a “competitor strain” that constitutively expresses blue fluorescent protein (BFP) (Fig. 6-a,b,c). The performance metric that we use for this comparison is the temporal population-level expression of RFP after its activation, that is, the intensity of RFP normalized by the OD of the co-culture. When grown in isolation and post induction of RFP, the growth rates of the OL and CL strains are initially close to each other and to that of the competitor strain. However, as time progresses, the growth rate of the OL strain drops to about 50% of its original value while that of the CL strain maintains the initial growth rate over time (Fig. 6-d).

**Figure 6:**
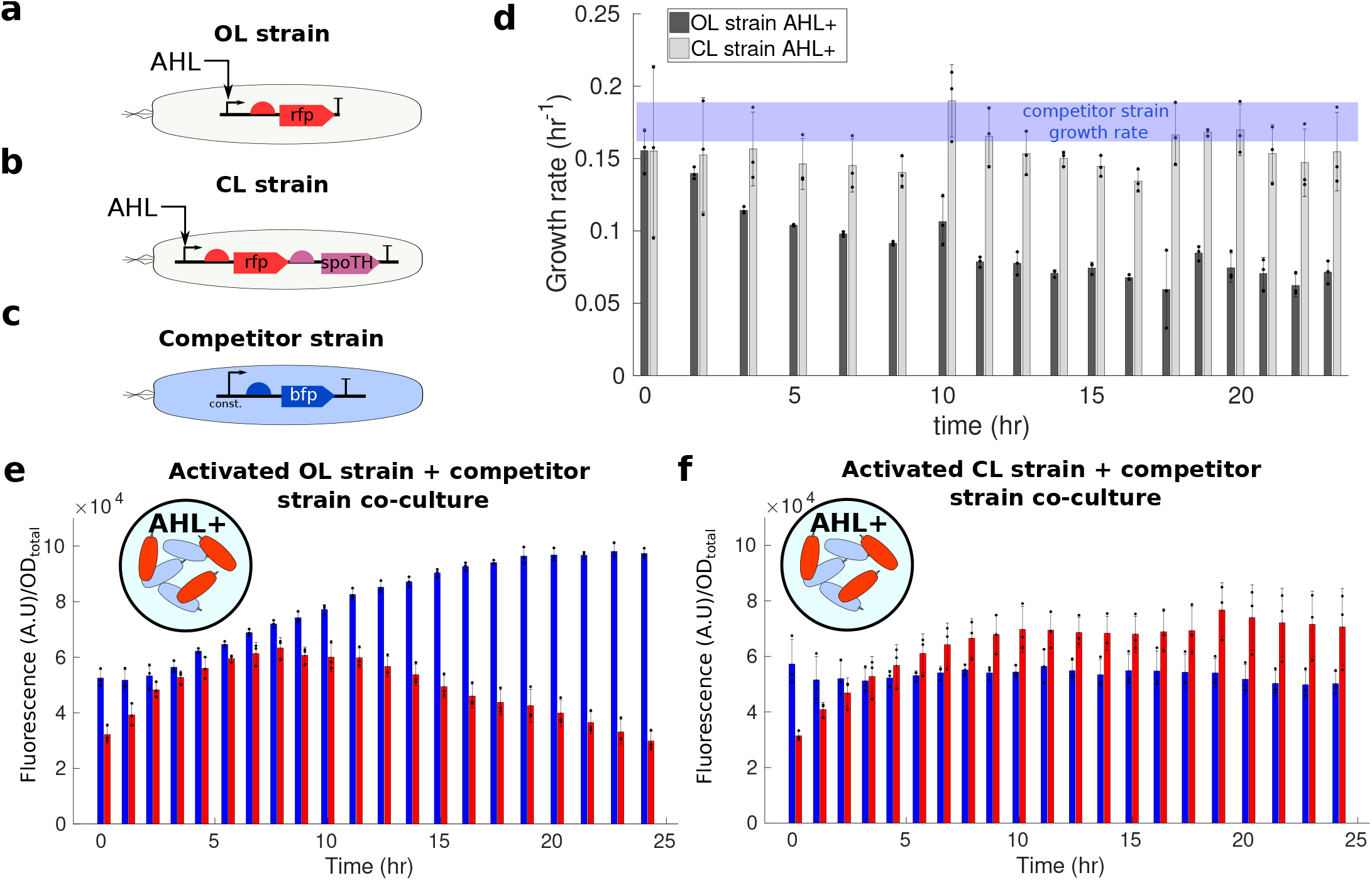
Feedforward controller promotes persistent GOI expression in a co-culture with a competitor strain. **(a)** The OL strain consists of P_OL in CF945. This strain expresses RFP using the AHL inducible Plux promoter. **(b)** The CL strain consists of P_IFFL___2 in CF945. This strain co-expresses RFP and SpoTH using the AHL inducible Plux promoter. **(c)** The competitor strain has P_BFP in TOP10. This strain constitutively expresses BFP. **(d)** Temporal responses of growth rate for the OL and CL strains grown in isolation post activation of the RFP gene (GOI) through AHL induction. The growth rate of the competitor strain grown in isolation is shown with a blue line (see SI Fig. 14 for raw data). **(e)-(f)** Temporal responses of RFP (red) and BFP (blue) fluorescence normalized by the total OD of the co-culture (OD_total_) for the OL and competitor strains co-culture in (e), and for the CL and competitor strains co-culture in (f). AHL+ denotes media containing the AHL inducer at 27.5 nM concentration. The growth rate and fluorescence of each strain for all biological replicates was simultaneously tracked in isolation (SI Fig. 14). Data are shown as the mean ± one standard deviation (N=3, three biological replicates). Individual experimental values are presented as black dots. All experiments were performed in media with glycerol as the sole carbon source. The complete experimental protocol is provided in the Materials and Methods section. Plasmid description, plasmid map, and essential DNA sequences are provided in SI section *Plasmid maps and DNA sequences*.

As a consequence, when OL and competitor strains are in co-culture and the GOI is activated, the population-level intensity of BFP increases, while that of RFP ultimately decreases (Fig. 6-e). This dynamic change in the population-level intensity of RFP and BFP can be attributed to the competitor strain overtaking the population due to its higher growth rate (compare blue line to dark gray bars in Fig. 6-d). To further verify that this population-level dynamic change was not due to a dynamic change of the expression level of BFP and RFP, we tracked the same biological replicates as in Fig. 6 in monoculture, which showed constant BFP and RFP intensity throughout the time course (SI Fig. 14). When the CL and competitor strains are in co-culture and the GOI is activated, the population-level intensity of BFP and RFP settle to a constant level (Fig. 6-f), consistent with the adaptation of the growth rate of the CL strain to its initial value post induction of the GOI (Fig. 6-d, light gray bars). Therefore, we conclude that the CL strain, by preventing a steady decrease in growth rate upon activation of the GOI, also allows persistent GOI population level expression.

## Discussion

The alarmone ppGpp has been referred to as the “CEO of the cell”, whose job is to optimally regulate resources for growth based on environmental conditions and current translational activity [52]. In this paper, we exploited the inverse relationship between ppGpp level and rRNA transcription rate during exponential growth [35, 36, 37, 38] and the hydrolysis of ppGpp by SpoT [42] to engineer an actuator that upregulates growth rate (Fig. 1). Specifically, the actuator exogenously expresses a modified version of spoT with only hydrolysis activity (SpoTH) and, in strains with elevated basal ppGpp level, activation of the SpoTH gene uprelgulates ribosomes (Fig. 2). We demonstrated the ability to actuate growth rate first in strains with elevated basal ppGpp level and by tuning the carbon source in the growth media (Fig. 2). Other methods such as tuning the amino acid concentration in the media could also be considered [35]. We then made the actuator portable to common laboratory strains by artificially raising ppGpp’s level through expression of the RelA+ enzyme (Fig. 4 and Fig. 5).

We employed the actuator to create a feedforward controller of ribosome level that compensates for the burden on cellular resources observed in the form of growth rate defects due to activating a GOI (Fig. 3 and Fig. 5). The controller co-expresses SpoTH with the GOI (RFP); therefore, when the GOI is activated, SpoTH is also activated, which increases ribosome availability. This increase in ribosome availability, when the SpoTH RBS is well tuned, exactly compensates for the GOI’s resource demand, leading to no change in growth rate (Fig. 3). The feedforward controller can be implemented for any GOI by co-expressing SpoTH with it and by tuning the SpoTH RBS based on the GOI’s ribosome load on the cell; hence, this design is tunable and modular.

For sufficiently low growth rates (high levels of ppGpp), the feedforward controller can also be tuned to keep the production rate of any constitutively expressed protein constant as the GOI is activated (SI Fig. 9 and SI Fig. 13). However, the SpoTH RBS that keeps growth rate constant is not the same as the one that keeps protein production rate constant. For a protein whose decay rate is dominated by degradation, the steady state concentration per-cell is approximately given by the ratio between the production rate and the degradation rate [53]. Therefore, the SpoTH RBS that keeps protein’s production rate constant as a GOI is activated, also keeps the protein’s concentration per-cell constant. In future applications, the feedforward controller may be used synergistically with previously engineered controllers that maintain concentration per cell constant once a resource competitor (GOI) is activated, but cannot maintain growth rate constant [21, 22, 23, 25, 24]. In fact, the concurrent implementation of the SpoTH feedforward controller with these controllers will both maintain constant growth rate and for any protein of interest and a constant concentration per-cell, when a GOI is activated.

The SpoTH actuator can also mitigate the growth defects caused by activation of a toxic protein, such as dCas9 [54, 55]. With the SpoTH actuator, we could reach without growth defects a dCas9 production rate that would otherwise cause a 40% decrease in growth rate without SpoTH expression (Supplementary Notes 7). We estimate that this production rate is at least four times higher than that reachable without growth defects without SpoTH expression (Supplementary Notes 7). These results have direct applications to CRISPRi-based genetic circuits where dCas9 should be at high concentrations to minimize the effects of its sequestration by multiple sgRNAs [56, 57, 58]. However, dCas9 toxicity limits its concentration to ranges where sequestration effects are prominent [56].

Persistent population-level expression of a GOI in a strain that shares the environment with competing organisms is hampered by growth rate imbalances that follow the GOI activation [4, 20]. We applied the feedforward controller to achieve persistent population-level expression of RFP (GOI) in a strain co-cultured with a competitor strain (Fig. 6). In applications, we can use the RelA+ inducible expression cassette to set the strain’s nominal growth rate to a desired level chosen to match the growth rate of other competitor strain(s), thereby achieving co-existence. The feedforward controller co-expressing SpoTH with the GOI then guarantees that this desired growth rate does not drop as the GOI is dynamically activated, thereby enabling persistent population-level expression of the GOI. This tool will thus be critical in future multi-strain systems that implement division of labor by running different genetic circuits with distinct, yet complementary, functionalities in the different strains [50, 51]. Population controllers have been implemented to promote coexistence in multi-strain systems. However, these controllers require the growth rates of each strain to be sufficiently close to one another [20]. In these systems, GOI activation in one strain may lower growth rate beyond the co-existence limits, at which point co-existence is lost despite the population controller. Our feedforward controller can be used synergistically with population controllers to ensure co-existence in multi-strain systems when expression of a GOI is dynamically modulated in each of the strains.

Our understanding of ppGpp, and especially of its role in exponential growth, is constantly evolving [52], so as we gain more insight on this pathway, we may uncover opportunities to further optimize the SpoTH actuator. For example, other enzymes like Mesh1 [59, 60] and SpoT E319Q [61] have been shown to catalyze the hydrolysis of ppGpp and can, in principle, serve in alternative actuator designs in place of SpoTH. Overall, this work provides an example of how we can exploit endogenous cell growth regulation pathways for engineering biology applications.

The feedforward controller is a tunable, modular, and portable tool that allows dynamic modulation of a GOI to possibly high-levels without substantially affecting growth rate. It will thus be a tool useful for all those applications where engineered bacteria need to co-exist with environmental species or with other engineered strains, while running circuits in which genes become dynamically activated.

## Materials and Methods

### Bacterial strain and growth

The bacterial strain used for genetic circuit construction was *E. coli* NEB10B (NEB, C3019I) and LB broth Lennox was the growth medium used during construction. Characterization was performed using the CF944, CF495, and CF946 [35], MG1655 (CGSC, 6300), and TOP10 strains. Characterization experiments were done using M9 minimal medium supplemented with 0.2% casamino acids,1 mM thiamine hydrochloride, ampicillin (100 *μ*g/mL), and either 0.4% glucose, 0.4% fructose, 0.4% glycerol, or 2 g/L lactose (the specific carbon source used for each experiment is specified in the figure caption).

### Microplate photometer protocol

This protocol was used to generate the data in all figures in the main text except that of the co-culture experiment (Fig 6). Cultures were prepared by streaking cells from a 15 % glycerol stock stored at − 80°C onto a LB (Lennox) agar plate containing 100 *μ*g/mL ampicillin and incubated at 37°C. Isolated colonies were picked and grown in 2 ml of growth medium in culture tubes (VWR, 60818-667) for 12-24 hours at 30°C and 220 rpm in an orbital shaker. Cultures were then diluted to an OD at 600 nm (OD_600nm_) of 0.0075 and grown for an additional 6 hours in culture tubes to ensure exponential growth before induction. Cultures were then induced and plated onto 96 well-plate (Falcon, 351172). The 96-well plate was incubated at 30°C in a Synergy MX (Biotek, Winooski, VT) microplate reader in static condition and was shaken at a fast speed for 3 s right before OD and fluorescence measurements. Sampling interval was 5 minutes. Excitation and emission wavelengths to monitor GFP fluorescence are 485 (bandwidth = 20 nm) and 513 nm (bandwidth = 20 nm), respectively and the Sensitivity = 80. Excitation and emission wavelengths to monitor RFP fluorescence are 584 (bandwidth = 13.5 nm) and 619 nm (bandwidth = 13.5 nm), respectively and the Sensitivity = 100. Sampling continued until bacterial cultures entered the stationary.

### Microplate photometer protocol for co-culture experiment

This protocol was used to generate the data for the co-culture experiment (Fig 6). Cultures were prepared by streaking cells from a 15 % glycerol stock stored at − 80°C onto a LB (Lennox) agar plate containing 100 *μ*g/mL ampicillin and incubated at 37°C. Isolated colonies were picked and grown in 2 ml of growth medium in culture tubes (VWR, 60818-667) for 12-24 hours at 30°C and 220 rpm in an orbital shaker. Cultures were then diluted to an OD at 600 nm (OD_600nm_) of 0.0075 for the OL and CL strain and 0.0045 for the competitor strain. After four hours the competitor strain was induced with 550 *μ*M SAL and after six hours the OL and CL strains were induced with 27.5 nM AHL. The cultures were then plated onto 96 well-plate (Falcon, 351172) and grown until the optical density was above OD_600nm_ = 0.02 and were then mixed to bring the optical density of the co-culture to OD_600nm_ = 0.02. The biological replicate of each culture was simultaneously tracked in isolation (mono-culutre). The cultures were then grown until one of the co-cultures reached OD_600nm_ = 0.2 and then all cultures were diluted to OD_600nm_ = 0.035, this dilution process was repeated three times (see SI Fig. 15 for growth curves). The 96-well plate was incubated at 30°C in a Synergy MX (Biotek, Winooski, VT) microplate reader in static condition and was shaken at a fast speed for 3 s right before OD and fluorescence measurements. Sampling interval was 5 minutes. Excitation and emission wavelengths to monitor BFP fluorescence are 400 (bandwidth = 9 nm) and 460 nm (bandwidth = 9 nm), respectively and the Sensitivity = 80. Excitation and emission wavelengths to monitor GFP fluorescence are 485 (bandwidth = 9 nm) and 513 nm (bandwidth = 9 nm), respectively and the Sensitivity = 80. Excitation and emission wavelengths to monitor RFP fluorescence are 584 (bandwidth = 13.5 nm) and 619 nm (bandwidth = 13.5 nm), respectively and the Sensitivity = 100.

### Calculating growth rate and protein production rates

The media background OD (0.08 OD_600nm_), GFP (100 A.U), and BFP (2,800 A.U) were subtracted from the data prior to any calculations. To ensure the data analyzed was coming from cells in exponential growth, only OD values (adjusted for background) of OD_600nm_ = 0.06 and OD_600nm_ = 0.14 were considered except for experiments done in lactose where the range was OD_600nm_ = 0.06 and OD_600nm_ = 0.1, since cells growing in lactose entered stationary phase at lower OD values.

To dampen noise before differentiating, the data was then filtered using a moving average filter. Given a signal with *n* measurements **y** = [*y*_1_, *y*_2_, …, *y*_*n*+1_] sampled at a constant period Δ*t*, we apply the moving average filter as follow:

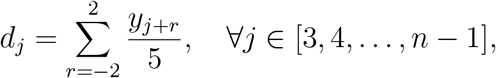

where **d** = [*d*_1_, *d*_2_, …, *d*_*n*+1_] is our filtered signal with boundary points identical to those of **y** (*d*_1_ = *y*_1_ and *d*_2_ = *y*_2_).

The growth rate is calculated from the filtered OD signal by performing linear regression (in a least-squares sense) on the log of the signal and taking the slope of the fit. The temporal growth rate data from Fig. 6-d was calculated by partitioning the OD versus time data (SI Fig. 15) into the time intervals shown in Fig. 6-d and then calculating the growth rate of each individual partition per the above method.

The RFP and GFP production rates were calculated in a similar manner as [62]. Denoting GFP(*t*_*i*_) and RFP(*t*_*i*_) as the filtered GFP and RFP signal measured by the plate reader at time *t*_*i*_, the GFP production rate (*α*_GFP_(*t*_*i*_)) and RFP production rate (*α*_RFP_(*t*_*i*_)) are given by

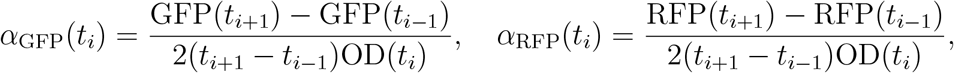

where OD(*t*_*i*_) is the filtered OD level.

### Genetic circuit construction

The genetic circuit construction was based on Gibson assembly [63]. DNA fragments to be assembled were amplified by PCR using Phusion High-Fidelity PCR Master Mix with GC Buffer (NEB, M0532S), purified with gel electrophoresis and Zymo clean Gel DNA Recovery Kit (Zymo Research,D4002), quantified with the nanophotometer (Implen, P330), and assembled withGibson assembly protocol using NEBuilder HiFi DNA Assembly Master Mix(NEB, E2621S). Assembled DNA was transformed into competent cells prepared by the CCMB80 buffer (TekNova, C3132). Plasmid DNA was prepared by the plasmid miniprep-classic kit (Zymo Research, D4015). DNA sequencing used Quintarabio DNA basic sequencing service. Primers and gBlocks were obtained from Integrated DNA Technologies. The list of constructs and essential DNA sequences can be found in SI section *Plasmid maps and DNA sequences*

## Acknowledgements

This work was supported in part by NSF Expeditions, Grant Number 1521925, NSF RoL Award, Grant Numbe 1840257, AFOSR grant FA9550-14-1-0060, the NSF Graduate Research Fellowships Program, and the Ford Foundation Predoctoral Fellowship. We thank Dr. Cashel, Dr. Potrykus, and Dr. Fernández-Coll for providing the CF944, CF945, and CF946 strains and their helpful discussion on ppGpp.

## Author contributions

D.D.V. and C.B designed the study; H.H, J.G, L.S, and C.B designed and built the genetic circuits; C.B performed the experiments; C.B analyzed the data; C.B developed the mathematical models; C.B and D.D.V. wrote the paper.

## Reporting summary

Further information on research design is available in the Nature Research Reporting Summary linked to this article.

## Data availability

Growth rate, fluorescence, and simulation data generated or analyzed during this study are included in the paper and its Supplementary Information files. A reporting summary for this article is available as a Supplementary Information file. Any other relevant data are available from the authors upon reasonable request.

## Code availability

Custom MATLAB (The MathWorks, Inc., Natick, MA, USA) codes are used to perform numerical simulations. A Supplementary Software file is provided.

## Supplementary Information

In the first section of the SI we provide supplementary experimental results. Then, we provide a detailed mathematical derivation of the SpoTH actuator model. Finally, we provide the plasmid maps of the constructs used in this study along with DNA sequence of nonstandard parts and end with supplementary notes.

### Additional experimental data

The GFP production rate data as SpoTH is expressed corresponding to Fig. 2 in the main text, is shown in Fig. 7. The RFP production rate vs growth rate data for all the CL RBS values tested for the experiment corresponding to Fig. 3 in the main text, is shown in Fig. 8. The GFP production rate data as the GOI is activated corresponding to Fig. 3 in the main text, is shown in Fig. 9. For the feedforward controller from Fig. 3-a in the main text, we replaced SpoTH with a nonfunctional heterologous protein CJB (cjBlue H197S [49]) and call this the control system. The growth rate and GFP production rate data as RFP is activated for the OL system and the control system is shown in Fig. 10. The GFP production rate data as RelA+ and SpoTH are expressed corresponding to Fig. 5 in the main text, is shown in Fig. 11. The RFP production rate vs growth rate data for all the CL RBS values tested for the experiment corresponding to Fig. 6 in the main text, is shown in Fig. 12. The GFP production rate data as the GOI is activated corresponding to Fig. 6 in the main text, is shown in Fig. 13. The fluorescence per-cell and growth rate of the same biological replicates as in Fig. 6 were simultaneously tracked individually in a mono-culture (Fig. 14). The growth curves for the mono-cultures and co-cultures corresponding to Fig. 6 in the main text are shown in Fig. 15.

**Figure 7:**
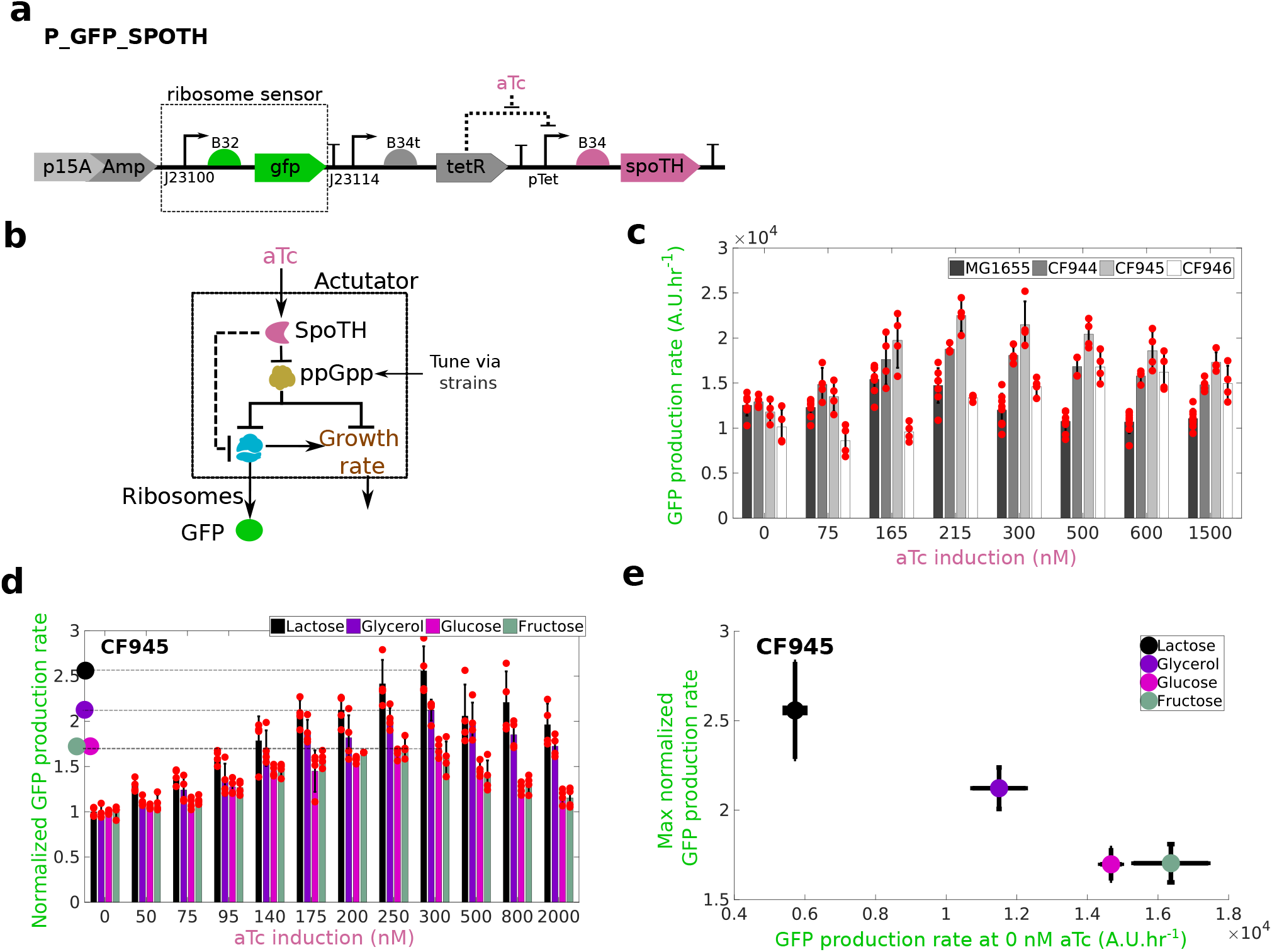
SpoTH expression increases GFP production rate. **(a)** The P_GFP_SpoTH plasmid used to express SpoTH via the inducible pTet promoter. Plasmid description, plasmid map, and essential DNA sequences are provided in SI section *Plasmid maps and DNA sequences*. **(b)** Addition of aTc increases SpoTH concentration, which lowers ppGpp concentration and consequently upregulates both free ribosome concentration and growth rate [35]. Here the constitutive GFP production rate serves as a proxy for free ribosomes (Supplementary note 4). **(c)** The GFP production rate while increasing SpoTH in the wild-type MG1655, CF944, CF945, and CF946 strains [35] growing in glycerol as the sole carbon source. **(d)** The GFP production rate normalized by the GFP production rate at aTc = 0 nM, as SpoTH is expressed in CF945 growing in lactose, glycerol, fructose, or glucose as the sole carbon source. The max normalized GFP production rate for each carbon source is marked by open squares. **(e)** The max normalized GFP production rate versus the GFP production rate at aTc = 0 nM for each carbon source. Data are shown as the mean ± one standard deviation (N=4, two biological replicates each with two technical replicates). Individual experimental values are presented as red dots. The complete experimental protocol is provided in the Materials and Methods section.

**Figure 8:**
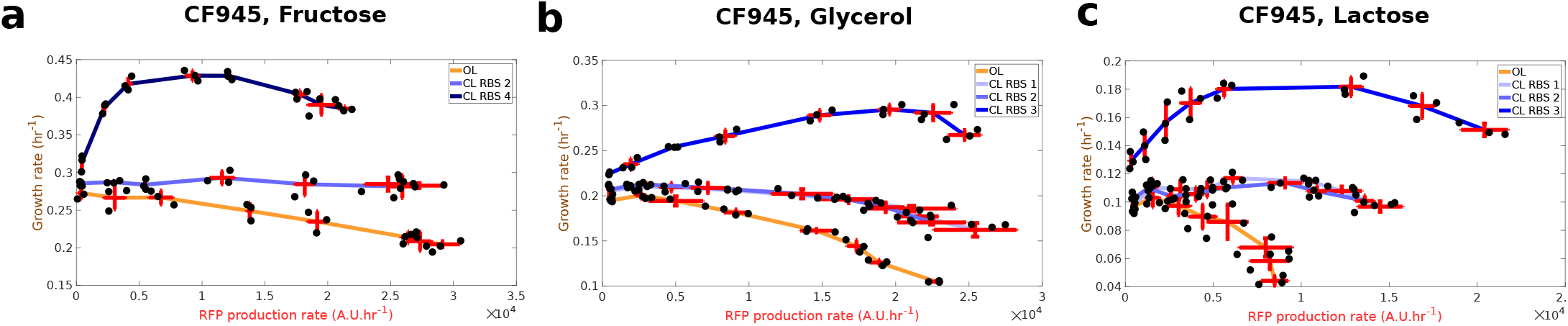
Feedforward ribosome controller compensates for burden caused by a GOI (RFP) activation. The unnormalized growth rate versus RFP production rate for all the tested RBS for the CL system corresponding to Fig. 3-e,f,g in the main text. **(a)-(c)** Growth rate versus the RFP production rate for the OL and CL systems, using fructose (e), glycerol (f), and lactose (g) as the carbon source. Data are shown as the mean ± one standard deviation (N=3, three biological replicates). All experiments were performed in the CF945 strain. Individual experimental values are presented as black dots. The complete experimental protocol is provided in the Materials and Methods section. Plasmid description, plasmid map, and essential DNA sequences are provided in SI section *Plasmid maps and DNA sequences*.

**Figure 9:**
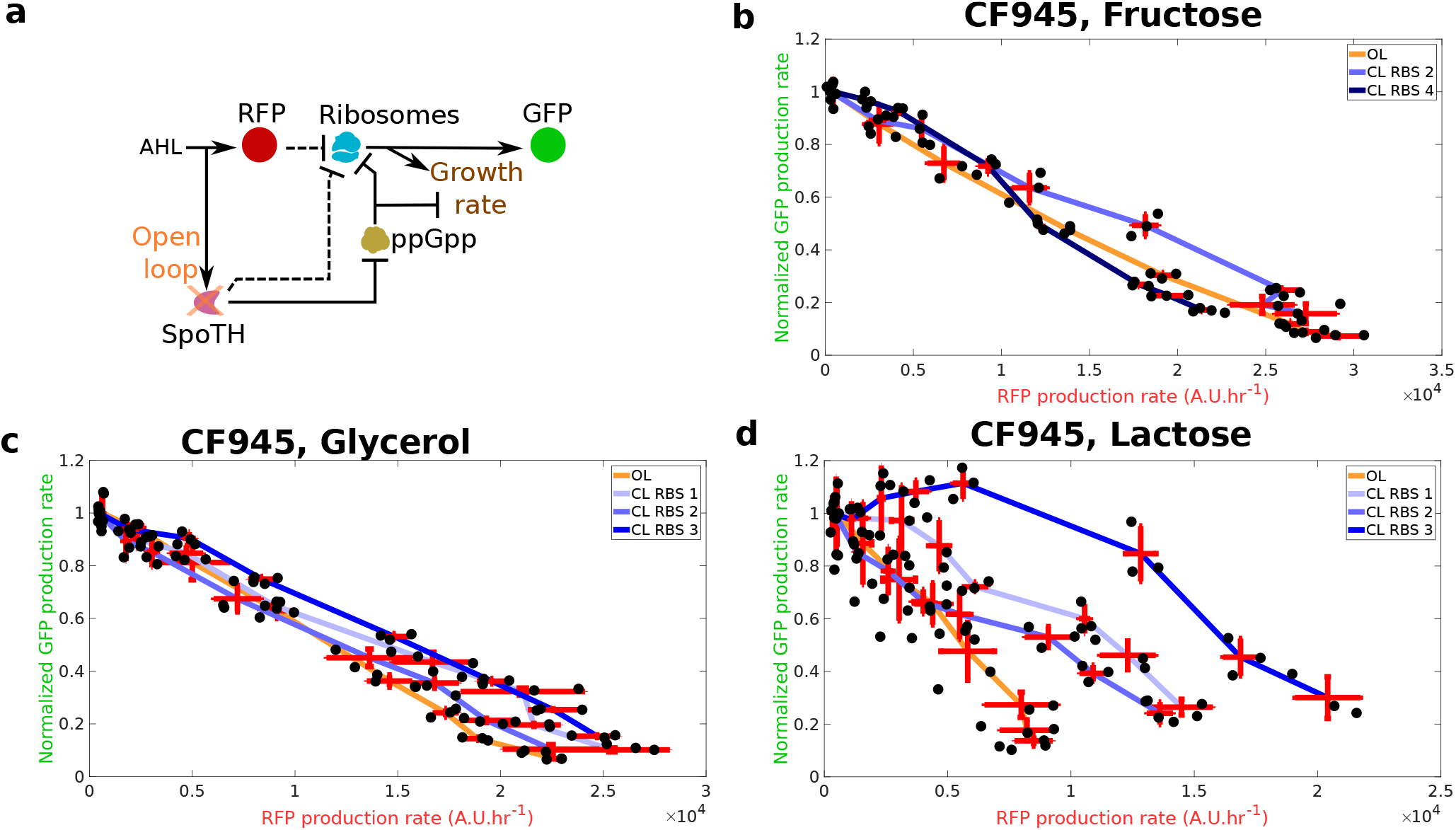
Feedforward controller compensates for GFP production rate defect caused by a GOI (RFP) activation at low growth rates. **(a)** Diagram depicting the effect of expressing RFP (via AHL) on ribosomes and growth rate for the open loop (OL) or closed loop (CL) systems. In the OL system, SpoTH is not present, so there is only the upper path from AHL to ribosomes. In the CL system, AHL also activates SpoTH expression and hence upregulates ribosome concentration and growth rate. Dashed edges represent sequestration of free ribosomes by a protein’s mRNA. Here the constitutive GFP production rate serves as a proxy for free ribosomes (Supplementary Note 4). **(b-d)** The GFP production rate normalized by the GFP production rate at 0 nM AHL versus the RFP production rate for the control, OL, and CL systems, using fructose (e) glycerol (f), and lactose (g) as the carbon source. Data are shown as the mean ± one standard deviation (N=3, three biological replicates). All experiments were performed in the CF945 strain. Individual experimental values are presented as black dots. The complete experimental protocol is provided in the Materials and Methods section. Plasmid description, plasmid map, and essential DNA sequences are provided in SI section *Plasmid maps and DNA sequences*.

**Figure 10:**
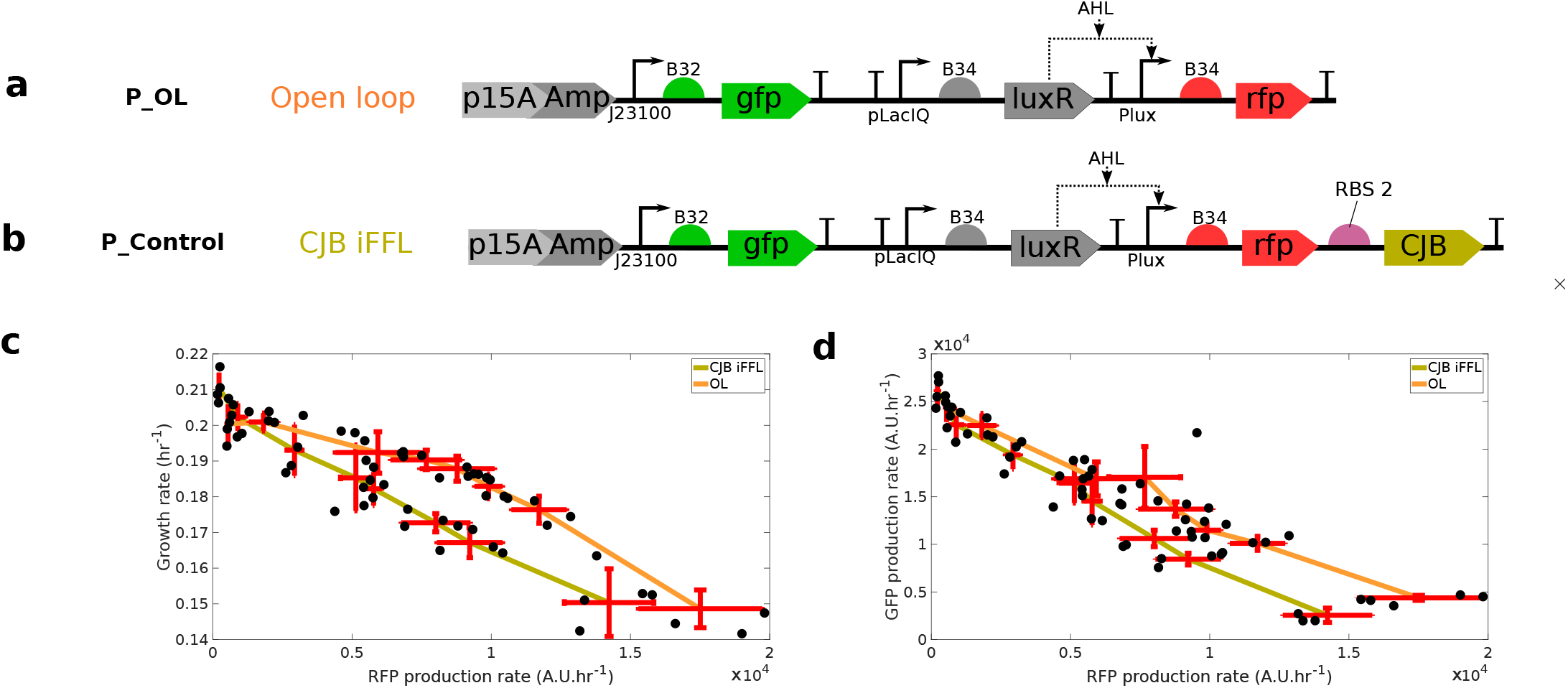
Replacing SpoTH with CJB in Feedforward controller creates more burden on growth rate and GFP production rate than the OL system. **(a)** OL system’s genetic construct (P_OL) used to express RFP using the AHL (TX input) inducible Plux promoter **(b)** The control genetic construct (P_Control) used to simultaneously express RFP and CJB. The CJB protein is a nonfunctional heterologous protein. **(c)/(d)** Growth rate/GFP production rate versus the RFP production rate for the control and OL systems. Data are shown as the mean ± one standard deviation (N=4, two biological replicates each with two technical replicates). Individual experimental values are presented as a black dots. All experiments were performed in the CF945 strain in media with glycerol as the sole carbon source. The complete experimental protocol is provided in the Materials and Methods section.

**Figure 11:**
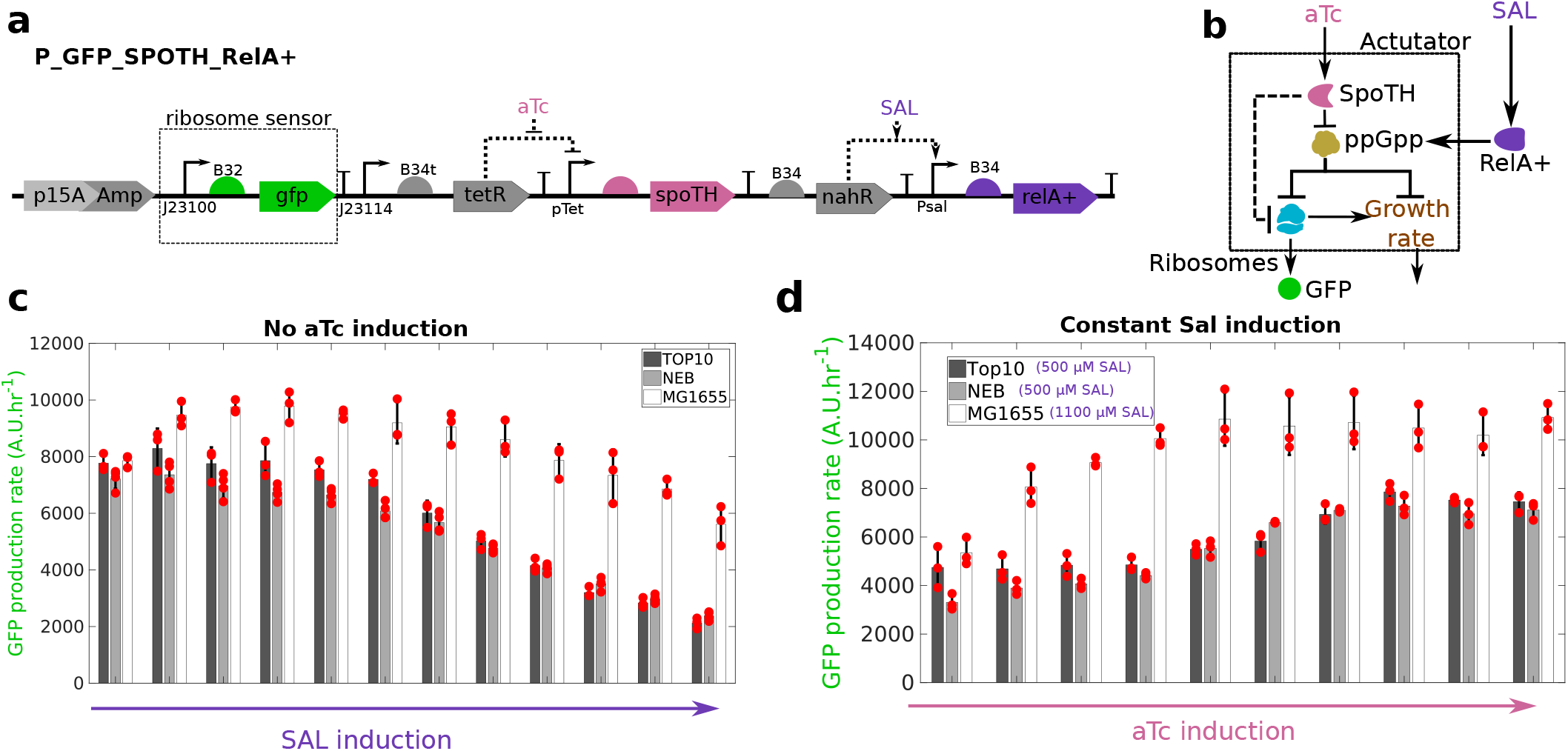
Expression of RelA+ allows to transport the SpoTH actuator to common laboratory strains. **(a)** The P_GFP_SpoTH_RelA+ plasmid used to express SpoTH via the inducible pTet promoter and RelA+ via the inducible Psal promoter. Plasmid description, plasmid map, and essential DNA sequences are provided in SI section *Plasmid maps and DNA sequences*. **(b)** Addition of SAL increases RelA+ concentration and thus upregulates ppGpp concentration [60]. Addition of aTc increases SpoTH concentration, which lowers ppGpp concentration and consequently upregulates both free ribosome concentration and growth rate [35]. Here the constitutive GFP production rate serves as a proxy for free ribosomes (Supplementary Note 4). **(c)** GFP production rate versus RelA+ induction (SAL) in the TOP10, NEB, and wild-type MG1655, strains growing in glycerol as the sole carbon source. **(d)** GFP production rate versus SpoTH expression (atc) for a fixed RelA+ expression in TOP10, NEB, and wild-type MG1655 strains growing in glycerol as the sole carbon source. Data are shown as the mean ± one standard deviation (N=3, three biological replicates). All experiments were performed with glycerol as the sole carbon source. Individual experimental values are presented as a red dots. The complete experimental protocol is provided in the Materials and Methods section.

**Figure 12:**
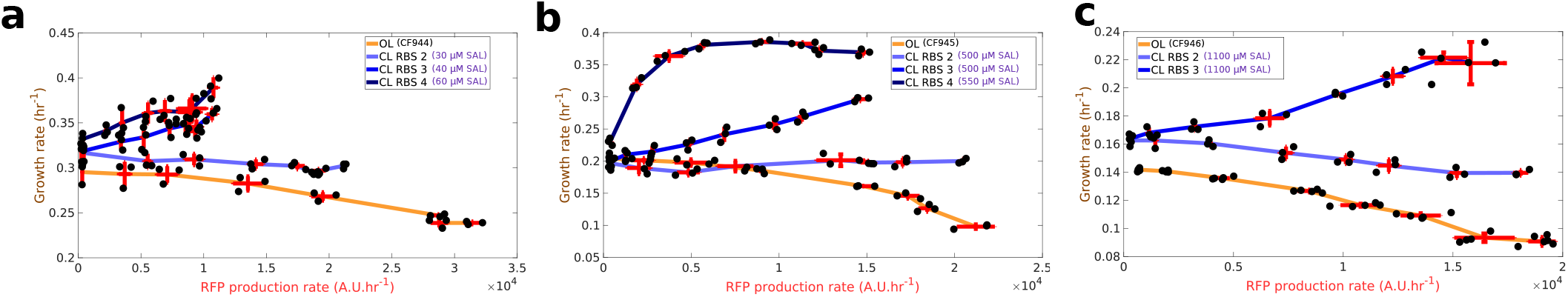
Feedforward ribosome controller compensates for burden caused by a GOI (RFP) activation in common bacterial strain. The unnormalized growth rate versus RFP production rate for all the tested RBS for the CL system corresponding to Fig. 6-c,d,e in the main text. **(a)-(c)** Growth rate versus the RFP production rate for the OL in CF944 (c), CF945, (d), and CF946 (e) and CL systems in TOP10. For the CL system, RelA+ expression is set to match the growth rate of the OL strain. All experiments were performed with glycerol as the sole carbon source. Data are shown as the mean ± one standard deviation (N=3, three biological replicates). All experiments were performed in the CF945 strain. Individual experimental values are presented as black dots. The complete experimental protocol is provided in the Materials and Methods section. Plasmid description, plasmid map, and essential DNA sequences are provided in SI section *Plasmid maps and DNA sequences*.

**Figure 13:**
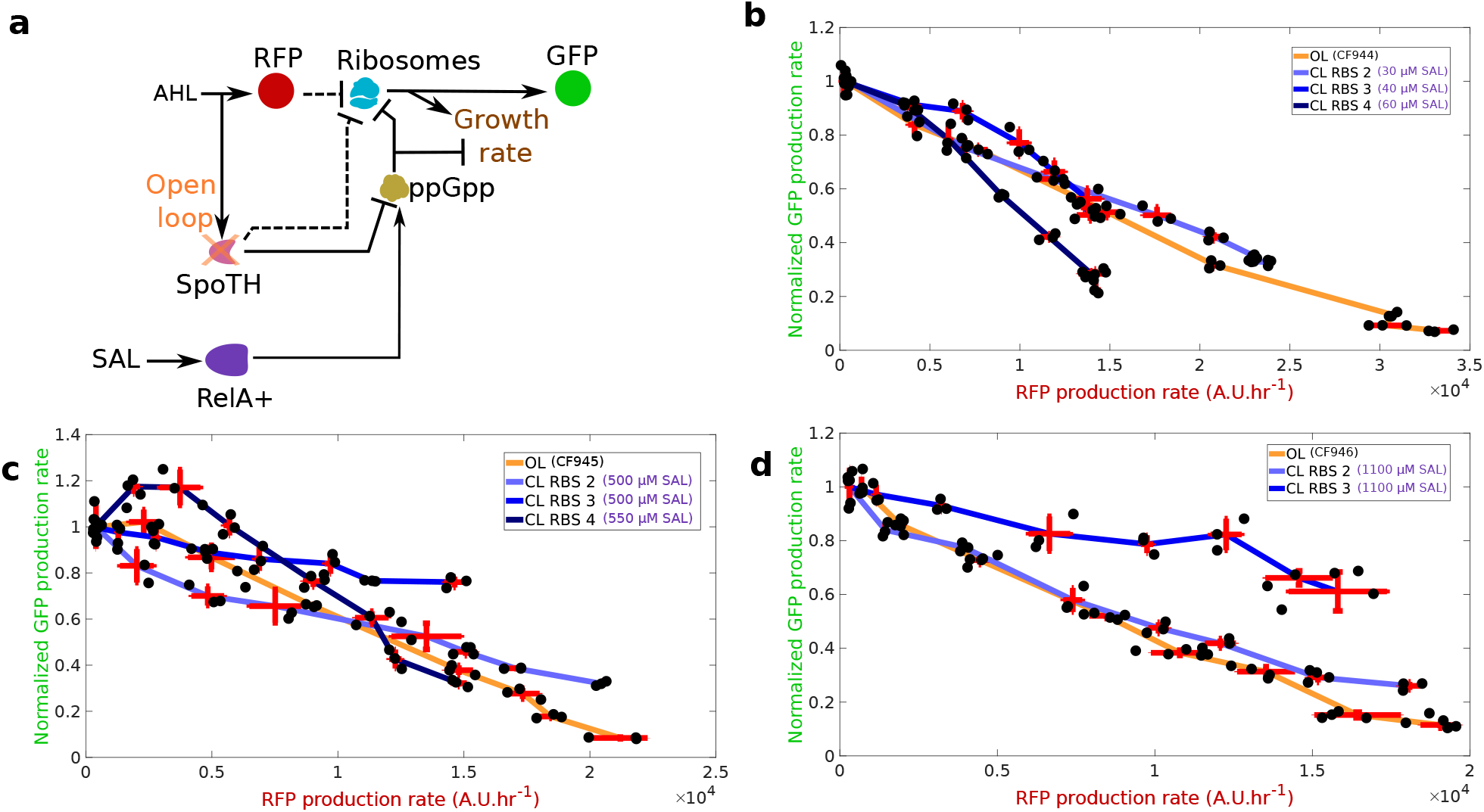
Feedforward controller compensates for GFP production defects caused by activation of RFP in TOP10 strain at low growth rates. **(a)** Diagram depicting the effect of activating RFP (AHL input) on ribosomes and growth rate for the open loop (OL) or closed loop (CL) systems. In the OL system, SpoTH is not present, so there is only the upper path from AHL to growth rate. In the CL system, the TX input also activates SpoTH production and hence upregulates ribosome concentration and growth rate. Dashed edges represent sequestration of free ribosomes by a protein mRNA. RelA+ activation via SAL sets the basal level of ppGpp and thus the setpoint growth rate [60]. Here the constitutive GFP production rate serves as a proxy for free ribosomes (Supplementary Note 4). **(c-e)** GFP production rate normalized by the GFP production rate at 0 nM AHL versus the RFP production rate for the OL in CF944 (c), CF945, (d), and CF946 (e) and CL systems (for RBS with best performance) in TOP10. For the CL system, RelA+ expression is set to match the growth rate of the OL strain. Data are shown as the mean ± one standard deviation (N=3, three biological replicates). All experiments were performed with glycerol as the sole carbon source. Individual experimental values are presented as black dots. The complete experimental protocol is provided in the Materials and Methods section. Plasmid description, plasmid map, and essential DNA sequences are provided in SI section *Plasmid maps and DNA sequences*.

**Figure 14:**
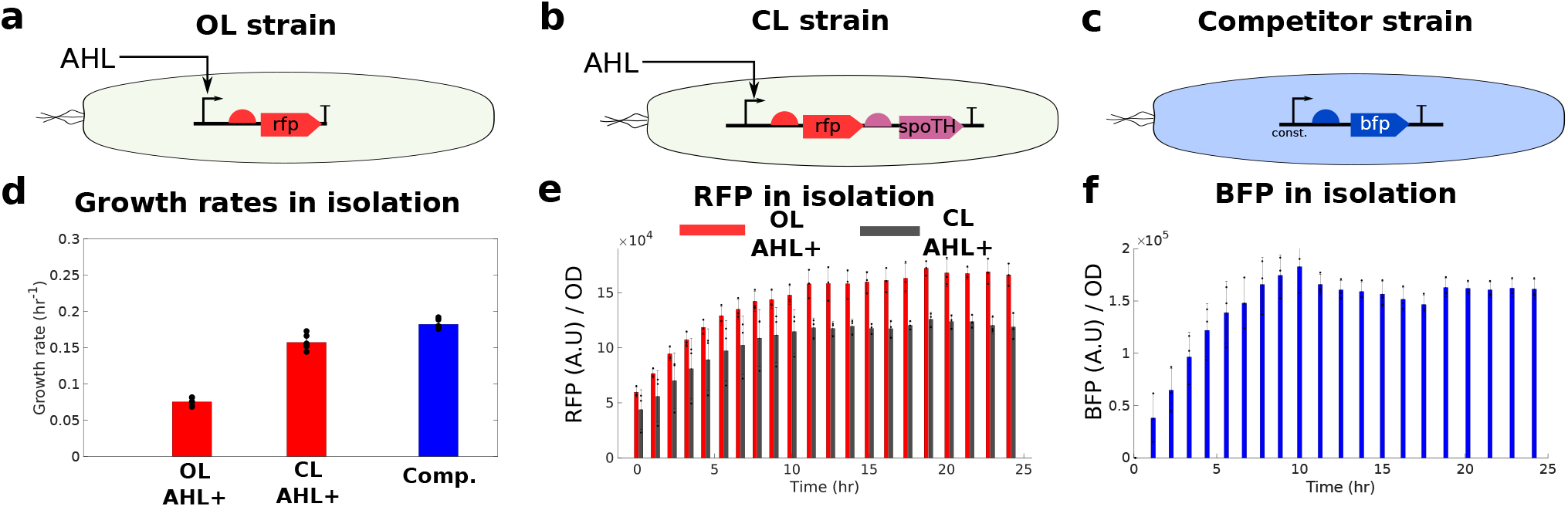
RFP and BFP expressed persistently in isolation. **(a)** The OL strain consists of P_OL in CF945. **(a)** The iFFL strain consists of P_IFFL___2 in CF945. **(c)** The competitor strain has P_BFP in TOP10. **(d)** The growth rates for each strain with AHL (AHL+, 27.5 nM) grown in isolation (mono-culture). The growth rates were calculated by averaging the growth rates of the last two batches in Fig. 15-a,b for each biological replicate. **(e)** The temporal response of mean RFP per OD values for the OL and CL strain with AHL (27.5 nM). **(f)** The temporal response of mean BFP per OD values for the competitor strain. Data are shown as the mean ± one standard deviation (N=3, three biological replicates). Individual experimental values are presented as a black dots. All experiments were performed in media with glycerol as the sole carbon source. Individual experimental values are presented as black dots. The complete experimental protocol is provided in the Materials and Methods section. Plasmid description, plasmid map, and essential DNA sequences are provided in SI section *Plasmid maps and DNA sequences*.

**Figure 15:**
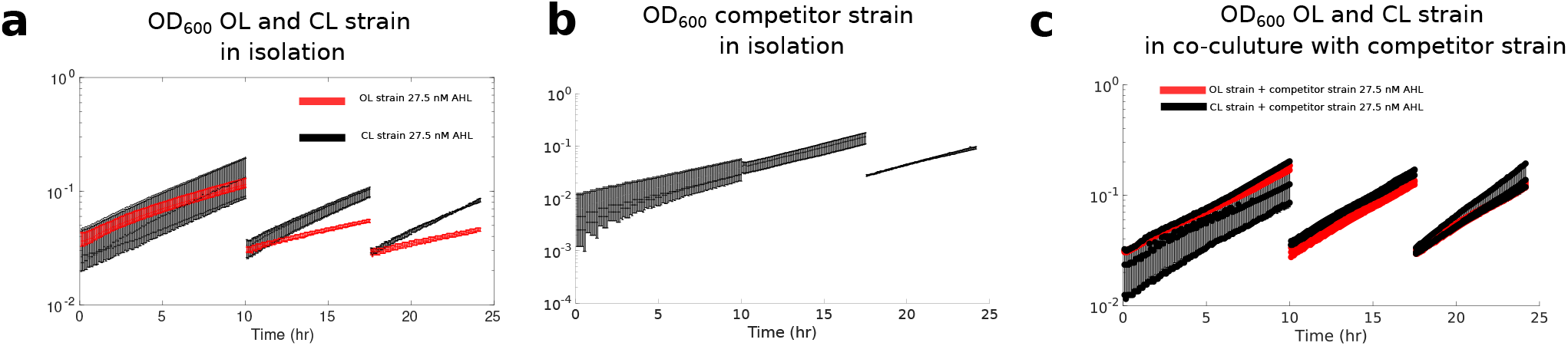
Co-culture experiment growth curves. **(a)** The optical density versus time for the OL and CL strain growing in mono-culture with AHL induction. **(b)** The optical density versus time for competitor strain growing in mono-culture. **(c)** The optical density versus time for the co-cultures of OL and competitor strain and the CL and competitor strain growing in co-culture with AHL induction. Data are shown as the mean ± one standard deviation (N=3, three biological replicates). Individual experimental values are presented as dots. All experiments were performed in media with glycerol as the sole carbon source. Individual experimental values are presented as black dots. The complete experimental protocol is provided in the Materials and Methods section. Plasmid description, plasmid map, and essential DNA sequences are provided in SI section *Plasmid maps and DNA sequences*.

### Derivation of the SpoTH actuator mathematical model

Following the deterministic modeling framework in [64] and previously applied in [13, 15], we derive a model of the SpoTH actuator. We model SpoTH mRNA being translated by ribosomes to produce the SpoTH protein, which catalyzes the hydrolysis of ppGpp. We model ppGpp inhibiting ribosome production and thus modifying the total ribosomal budget. The resulting dimensional model contains many free parameters, by nondimensionalizing the equations, we can reduce our governing equation to contain only two dimensionless parameters. Finally, we modify the equations to account for the expression of a heterologous protein.

This modeling framework is not meant to be comprehensive, but rather contain sufficient fidelity to make mathematically precise the physical processes discussed in the main text. The mathematical model is meant to complement the physical intuition provided in the main text used to explain the experimental data.

#### SpoTH expression and ppGpp hydrolysis and synthesis

We model SpoTH mRNA (m_s_) binding to free ribosomes (R) to produce the translation initiation complex c_s_, which is then translated to produce the SpoTH protein S with elongation rate constant *κ*_*s*_. The mRNA decays with rate constants *δ*_*s*_ and the protein dilutes with rate constant *γ*_*s*_. The corresponding chemical reactions are:

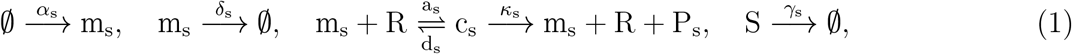

where *α*_*s*_ is the production rate constant of the mRNA, *a*_*s*_ and *d*_*s*_ are the association and dissociation rate constant, respectively, between ribosomes and mRNA. Levering reaction rate equations, consequently, the concentration of each species satisfies:

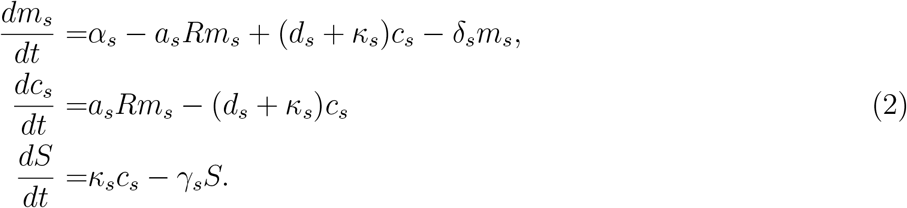

The steady state of (2) is given by

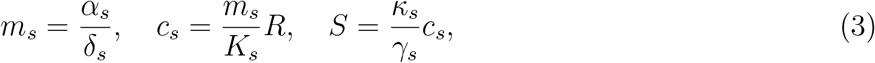

where 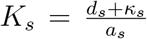. From (3), the concentration of SpoTH *S* is proportional to *c*_*s*_ (the number of ribosomes translating SpoTH mRNA).

We model RelA (A) and endogenous SpoT (S_0_) catalyzing the synthesis of ppGpp (G) (as in Fig. 1 in the main text) from GTP/GDP (G_P_), with rates *s*_1_ and *s*_2_, respectively. We model endogenous SpoT and SpoTH catalyzing the hydrolysis of ppGpp to GTP/GDP (G_P_) with rates *h*_1_ and *h*_2_, respectively. For simplicity, we model these processes using a *one-step reaction model* [64], that is,

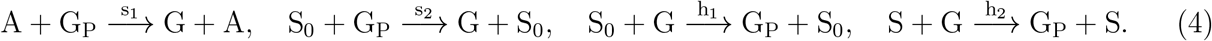

The concentration of ppGpp satisfies:

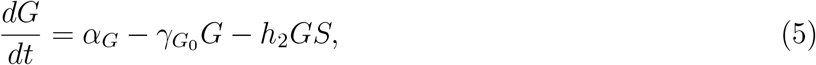

where *α*_*G*_ = *s*_1_*AG*_*P*_ + *s*_2_*S*_0_*G*_*P*_ is the effective production rate and 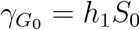 is the basal decay rate. The steady state of (5) is given by

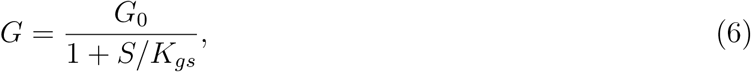

where 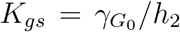 and 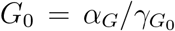. The quantity *G*_0_ corresponds to the basal ppGpp in the cell (*G*(*S* = 0) = *G*_0_). The quantity *G*_0_ is varied experimentally via chromosolmal mutation (e.g., *spot203*), media carbon source, and RelA+ expression (Fig. 2 and Fig. 5 in the main text).

#### Actuating the ribosomal budget in the cell

The concentration of total ribosomes in the cell (*R*_*T*_), known as the ribosomal budget [13], is composed of free ribosomes, the portion of ribosomes translating endogenous mRNAs (*c*_*e*_), and the portion of ribosomes translating SpoTH mRNA, that is,

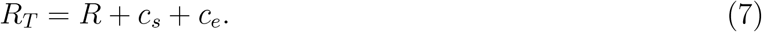

The total ribosome concentration obeys

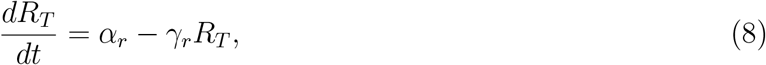

where *α*_*r*_ is the ribosome production rate, *γ*_*r*_ is the ribosome decay rate. If *α*_*r*_ and *γ*_*r*_ are assumed time invariant and 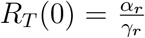, then 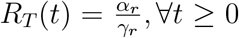. A temporally constant ribosome budget is consistent with the modeling framework of [13, 15, 22]. However, in this work *α*_*r*_ is not constant and is the term that links ribosome and ppGpp concentration.

Ribosome production (*α*_*r*_) is set by rRNA production (this the rate-limiting step) [30, 65]. rRNA is expressed from seven rRNA operons (*rrn* operons) [66] each driven by two tandem promoters P1 and P2. Most rRNA transcription arises from the P1 promoter and it is the main “knob” for ribosome tuning except at very low growth rates where P2 regulation dominates [67]. During balanced exponential growth, ppGpp is the primary regulator of rRNA[30, 31, 32] by destabilizing the open RNAP-P1 promoter complex [68, 69]. Therefore, there is an inverse relationship between basal ppGpp levels and rRNA transcription [35, 36, 37, 38]. A simple model to capture this process (previously used in [11]), is given by

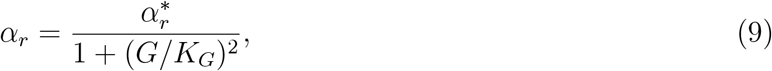

where *K*_*G*_ is the effective dissociation constant between ppGpp and the P1 promoter and 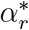 is the ribosome production rate in the absence of ppGpp (*α*_*r*_(*G* = 0)). The hill coefficient of 2 in (9) is consistent with the findings of [66]. Taking the steady state of (8) and levering (3), (6), and (9), we have that

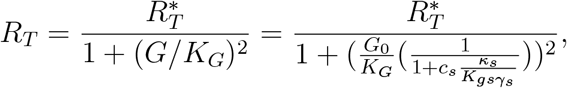

where 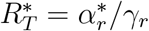.

Rewriting (7), we have at steady state that

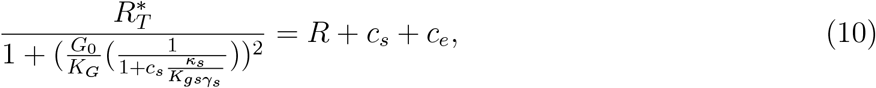

making explicit the relationship between basal ppGpp concentration (*G*_0_) and the total ribosomal budget and how increasing SpoTH expression (increasing *c*_*s*_) both increases the total ribosomal budget (LHS) but also sequesters ribosomes (RHS) via translation demand.

By adding and subtracting 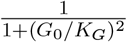 to the LHS of (7) and dividing both sides by 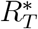, we can rewrite of (7) as

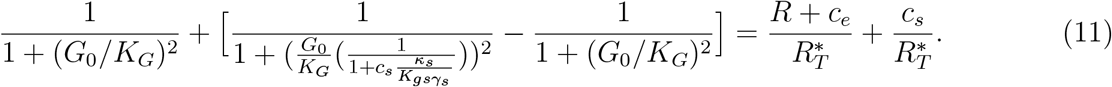

Modeling *c*_*e*_ requires knowing the concentration of the mRNA-ribosome complex for every mRNA expressed by an endogenous gene and thus it is difficult to write an explicit expression. Instead of modeling *c*_*e*_ explicitly, we keep it as a general function of *R*. We assume that the concentration of *c*_*e*_(*R*) monotonically increases with free ribosomes, that is, 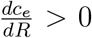 This assumptions is reasonable since a steady state complex concentration is proportional to the concentrations of the reacting species [53] (also (3)). Next, we define the following variable that serves as a proxy for free ribosomes:

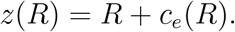

From the assumption that 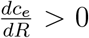, it implies that the map *z*(*R*) is one-one and thus for every value of *R* there is a unique corresponding value of *z* and that an increase/decrease in *R* corresponds to an increase/decrease in *z*. Furthermore, we have that *z*(0) = 0 since no the complex *c*_*e*_ cannot be formed without the reactant species *R*. Therefore, from here on, we refer to *z*(*R*) as the *modified free ribosome concentration*.

*Example:* if we assume that *c*_*e*_(*R*) had a form similar to that as *c*_*s*_ as given by (3), then, for *q* different endogenous genes expressing mRNA, 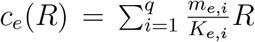, where for gene *i, m*_*e,i*_ is the endogenous mRNA concentration and *K*_*e,i*_ is the effective dissociation constant of endogenous mRNA with ribosomes. In this cases, *c*_*e*_(*R*) satisfies all of our assumptions and furthermore, *z*(*R*) is simply proportional to *R*. However, in all of our analysis we do not explicitly specify *c*_*e*_(*R*).

We denote 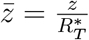 and 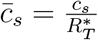 and express (11) in dimensionless form as

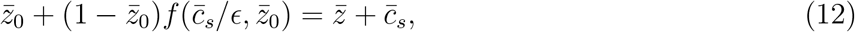

where

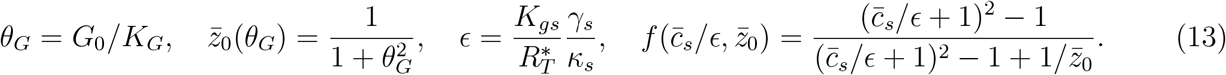

The dimensionless parameter *θ*_*G*_ is a measure of the basal ppGpp in the cell, 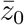 is the dimensionless modified free ribosome concentration when no SpoTH is expressed 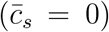 and we refer to this quantity as the *nominal modified ribosome level, ϵ* is a measure of the ribosomal cost to express sufficient SpoTH to actuate (catalyze the hydrolysis of a sufficient amount of ppGpp). A small *ϵ* implies that a small 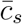 is needed to saturate the *f* term. Also notice that there is a monotonically decreasing relationship between the basal ppGpp *θ*_*G*_ and the nominal modified ribosome level. Finally, a key parameter to determine the qualitative behavior of (12) is given by:

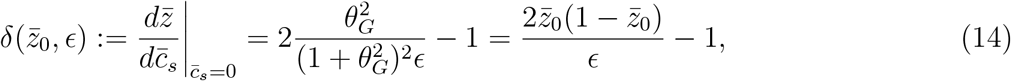

where *δ* ∈ (−1, ∞). By definition and our assumption that 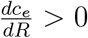, if *δ* > 0, it implies that ribosome levels increase as a small amount of exogenous SpoT is expressed.

#### Appending the model with the expression of an additional heterologous protein

We model the mRNA of a heterologous protein (m_y_) binding to free ribosomes (R) to produce the translation initiation complex c_y_, which is then translated to produce the protein y with elongation rate constant *κ*_*y*_. The mRNA decays with rate constants *δ*_*y*_ and the protein dilutes with rate constant *γ*_*y*_. The corresponding chemical reactions are:

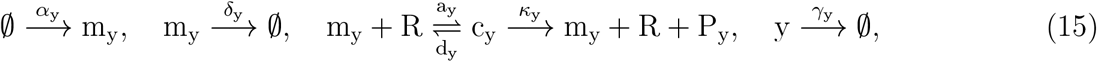

where *α*_*y*_ is the production rate constant of the mRNA, *a*_*y*_ and *d*_*y*_ are the association and dissociation rate constant, respectively, between ribosomes and mRNA. The concentration of each species satisfies:

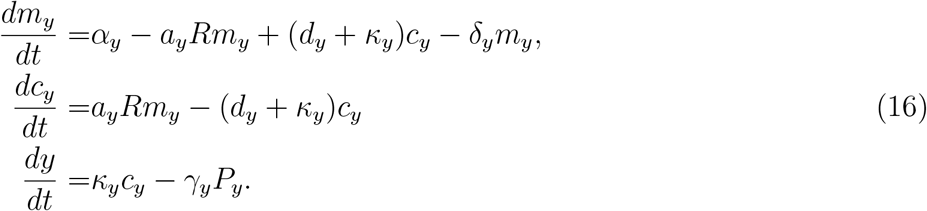

The steady state of (16) is given by

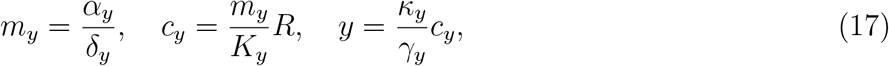

where 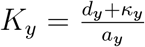. We modify the total ribosome equation (7) to include the ribosomes sequestered by the y mRNA, and it reads

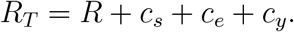

Defining 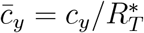, the total ribosome concentration in dimensionless from as in (12), is given by

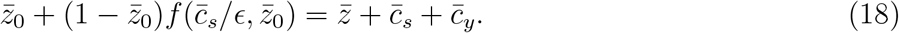

If y and SpoTH are under the same promoter, that is *m*_*y*_ = *m*_*s*_, then from (3) and (17) we have that at steady state

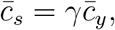

where *γ* = *K*_*y*_*/K*_*s*_ is the SpoTH RBS strength relative to the y RBS strength. We refer to the configuration when SpoTH and y are under the same promoter 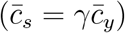, as the closed loop and the case when y is expressed in isolation (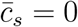 for all 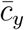), as the open loop. The qualitative behavior of (18) for the open loop and closed loop is shown in Fig. 16. For the close loop, we have can express the initial sensitivity of free ribosome as y is expressed as

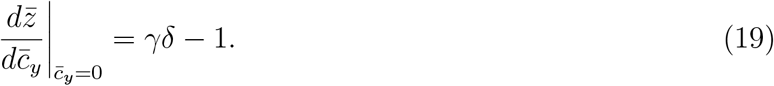

Thus, we can make the slope zero (free ribosomes are initially not sensitive to the expression of y) if we choose the SpoTH RBS strength (relative to the y RBS strength) as

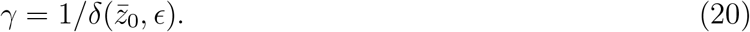

In Fig. 16 we observe that as *γδ* → − 1, the closed loop performs worst than the open loop (for a given 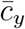, the corresponding value of 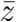 is lower) and the performance gets worst for larger values of *γ*.

### Plasmid maps and DNA sequences

The plasmids used in this study and their description are provided in Table 1. The corresponding plasmid maps are shown in Fig. 22. The essential DNA sequences are provided in Table 2. The full plasmid DNA sequences are uploaded to Addgene (#*xxx* − *xx*, will specify once study is finalized).

**Table 1:** Description of plasmids used in this study

| Plasmid Name | Plasmid map | Comments |
| --- | --- | --- |
| P_GFP_SpoTH | Fig. 22-a | SpoTH cloned using pHX41 [43], see Supplementary Note 3<br>The rest of the parts are from pHH03C_32 [22] |
| P_GFP_SpoTH_RFP | Fig. 22-b | P_GFP_SpoTH with luxR-Plux-RFP added from MBP-1.0 [13] |
| P_GFP_SpoTH_dCas9 | Fig. 22-c | P_GFP_SpoTH_RFP with RFP replaced by dCas9 from pdCas9_OP in [58] |
| P_OL | Fig. 22-d | P_GFP_SpoTH_RFP with TetR-pTet-SpoTH removed |
| P_IFFL_1 | Fig. 22-e | P_OL with RBS 1-SpoTH directly downstream of RFP |
| P_IFFL_2 | Fig. 22-e | P_OL with RBS 2-SpoTH directly downstream of RFP |
| P_IFFL_3 | Fig. 22-e | P_OL with RBS 3-SpoTH directly downstream of RFP |
| P_IFFL_4 | Fig. 22-e | P_OL with RBS 4-SpoTH directly downstream of RFP |
| P_Control | Fig. 22-f | P_IFFL_2 but SpoTH replaced by CJB (cjBlue H197S [49])<br>The CJB DNA is codon optimized for *E. coli* |
| P_weak_RFP | Fig. 22-d | P_OL but changed RFP RBS from B34 to RBS weak (MBP-0.006 in [13]) |
| P_weak_RFP_SpoTH_1 | Fig. 22-e | P_IFFL_1 but replaced RFP RBS from B34 to RBS weak |
| P_weak_RFP_SpoTH_2 | Fig. 22-e | P_IFFL_2 but changed RFP RBS from B34 to RBS weak |
| P_weak_RFP_SpoTH_3 | Fig. 22-e | P_IFFL_3 but changed RFP RBS from B34 to RBS weak |
| P_weak_RFP_SpoTH_4 | Fig. 22-e | P_IFFL_4 but changed RFP RBS from B34 to RBS weak |
| P_GFP_SpoTH_RelA+ | Fig. 22-g | P_GFP_SpoTH but with an inducible RelA+ [60] cassette |
| P_IFFL_SpoTH_RelA_1 | Fig. 22-h | P_IFFL_1 with an inducible RelA+ cassette |
| P_IFFL_SpoTH_RelA_2 | Fig. 22-h | P_IFFL_2 with an inducible RelA+ cassette |
| P_IFFL_SpoTH_RelA_3 | Fig. 22-h | P_IFFL_3 with an inducible RelA+ cassette |
| P_IFFL_SpoTH_RelA_4 | Fig. 22-h | P_IFFL_4 with an inducible RelA+ cassette |
| P_BFP | Fig. 22-i | inducible RelA+ cassette with constitutive BFP from [70] |

**Table 2:**
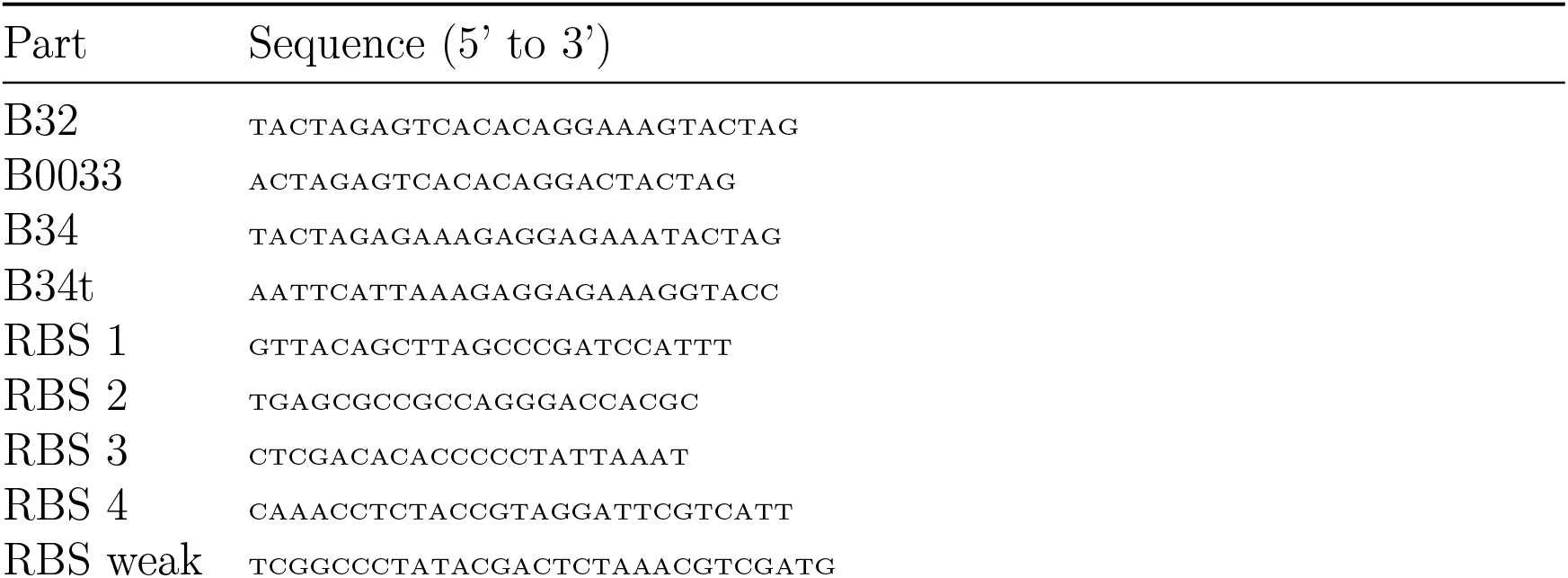
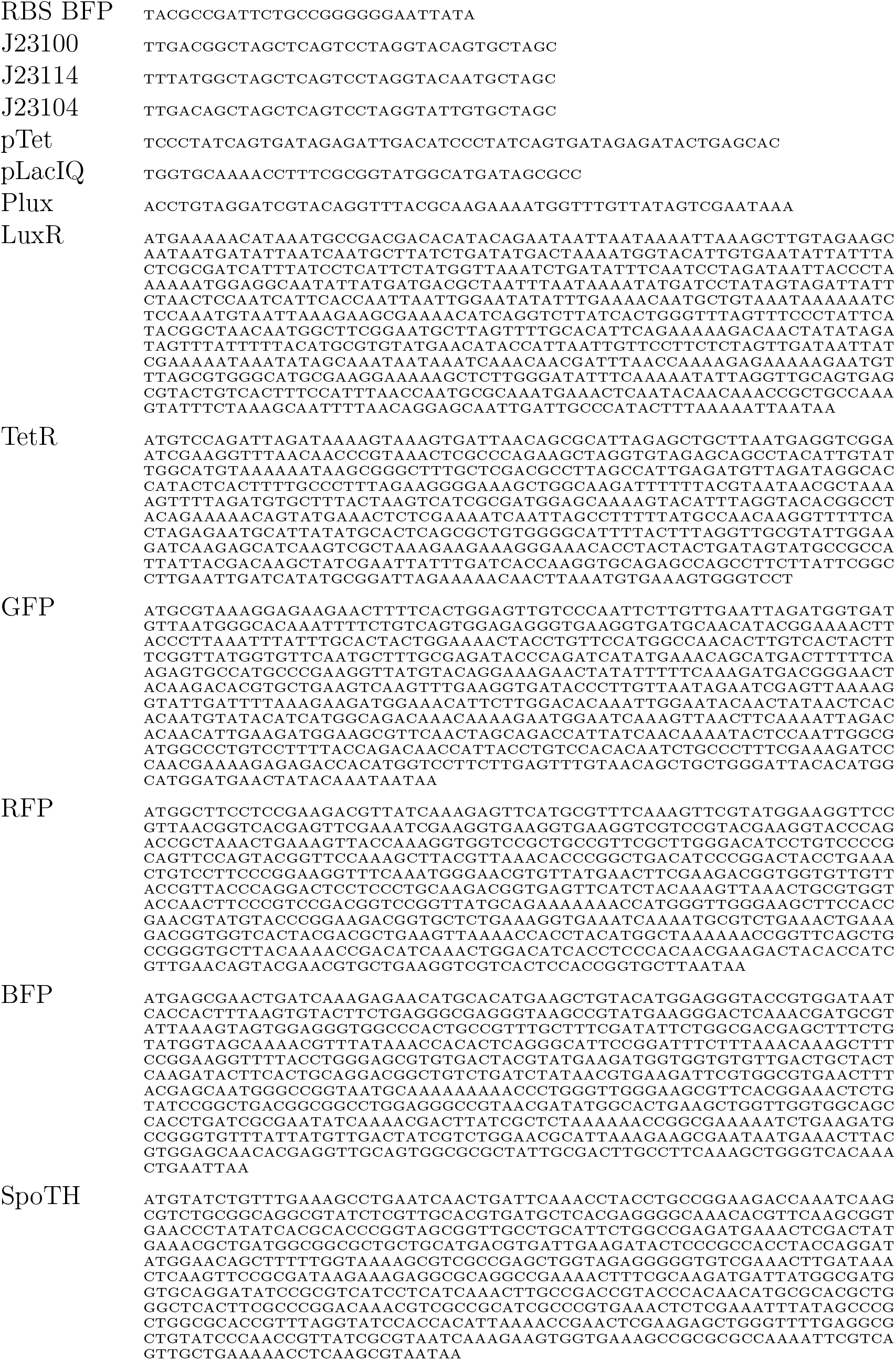
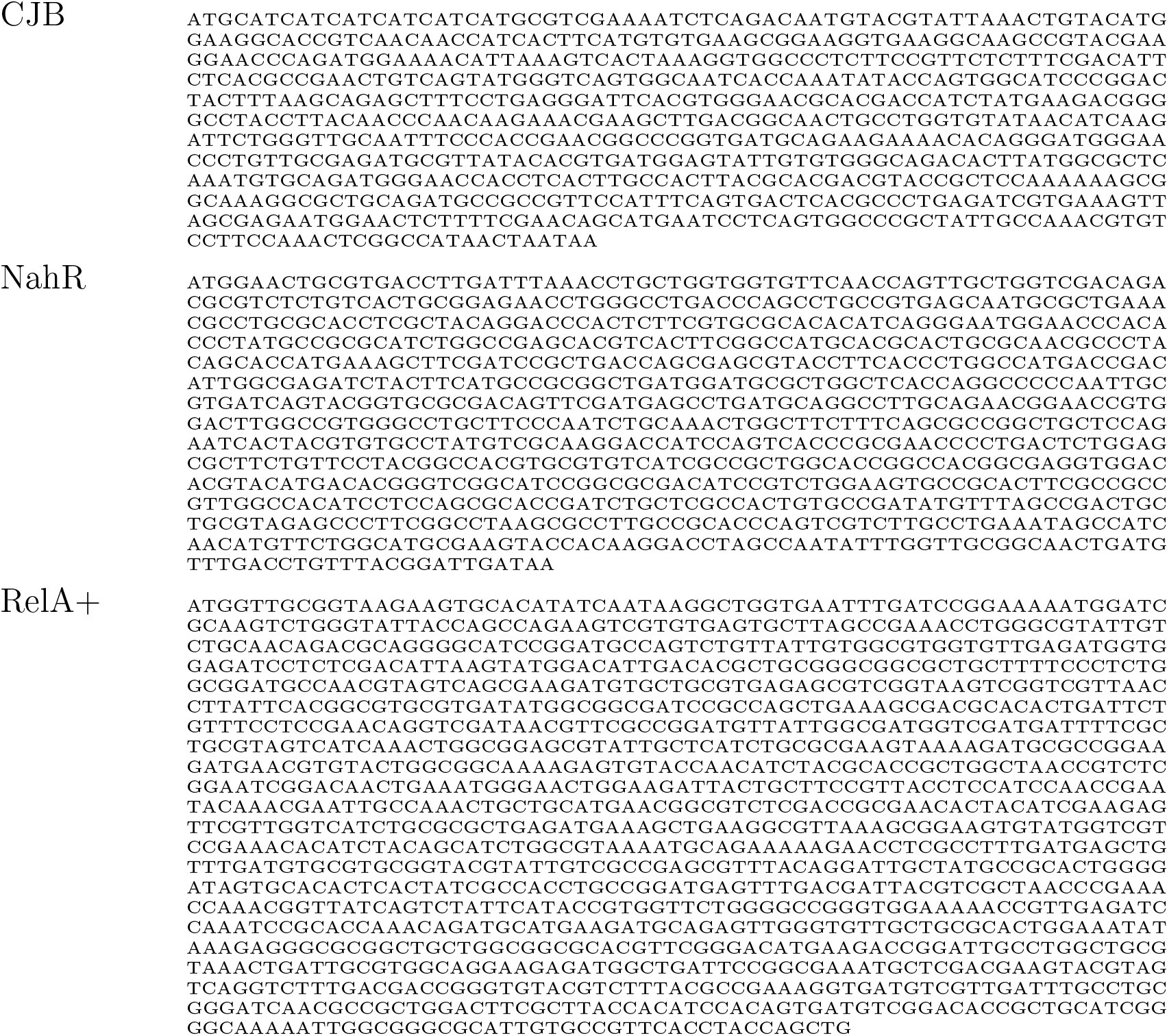
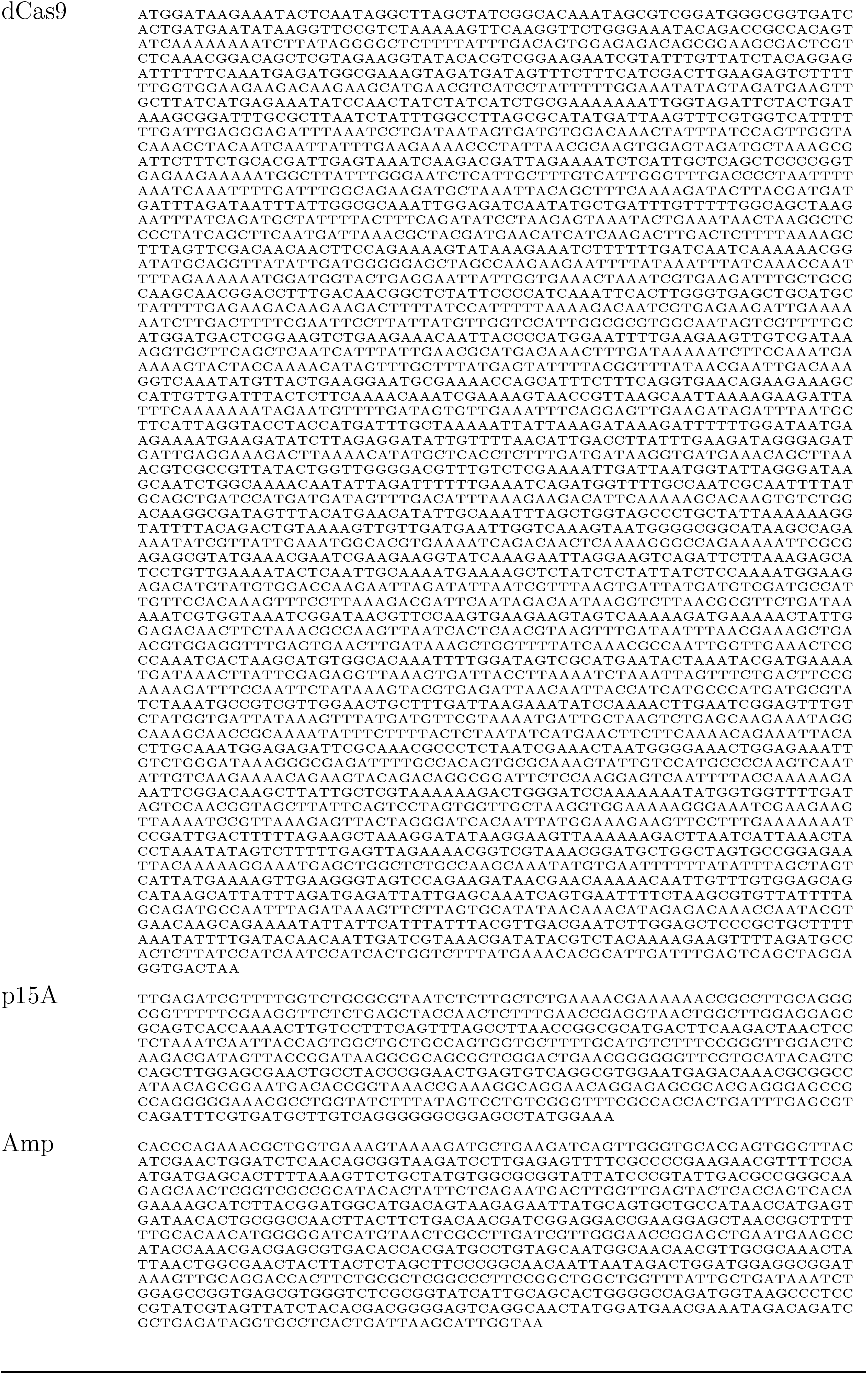
Essential DNA sequences used in this study.

**Figure 16:**
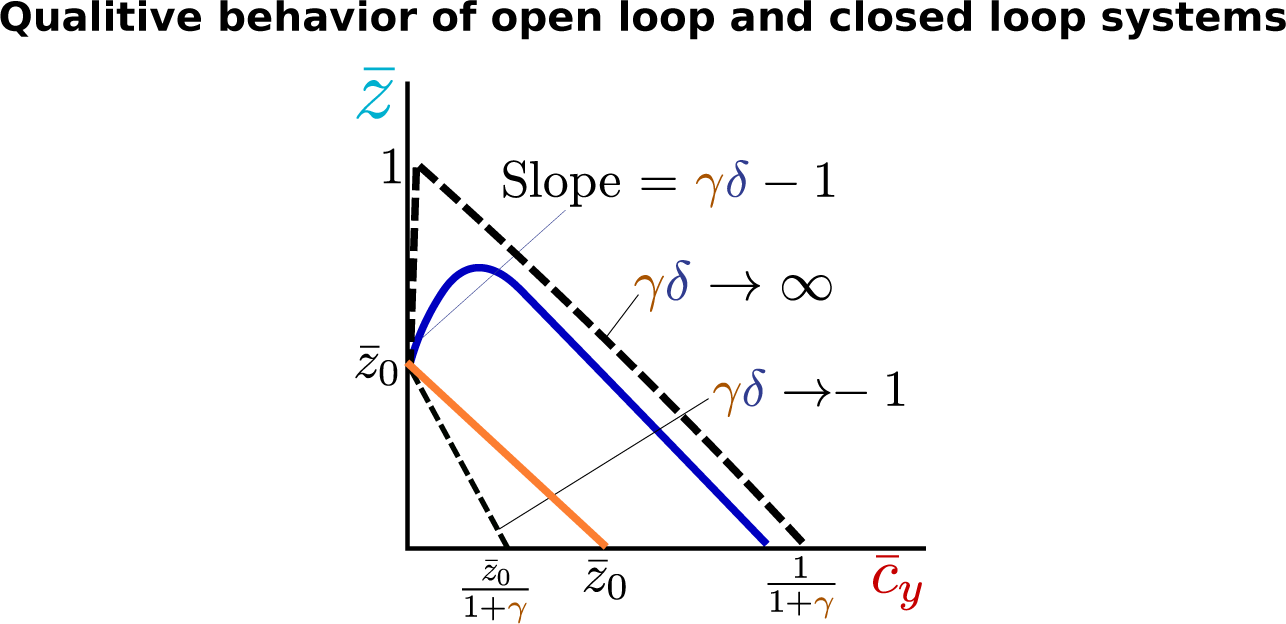
The qualitative behavior coexpressing y and SpoTH under the same promoter. The qualitative behavior of (18) when 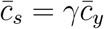 (blue line). The open loop (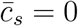 for all 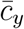) is shown in orange. The asymptotic behaviors as *γδ* → −1 and *γδ* → ∞ are shown in dashed lines.

**Figure 17:**
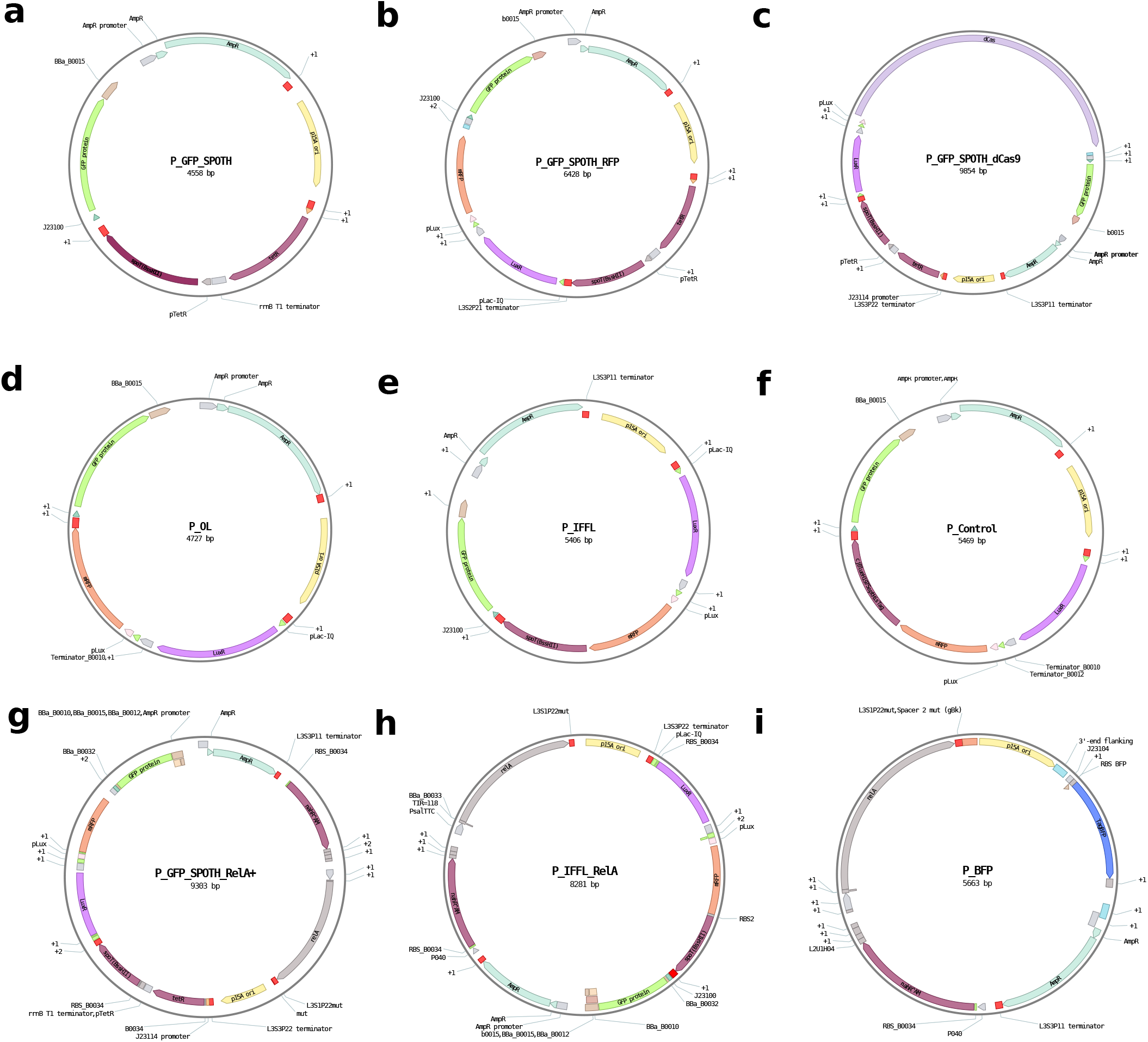
Plasmid maps. The plasmid maps were prepared by the Benchling Life Sciences R&D platform. **(a)** P_GFP_SpoTH **(b)** P_FP_SpoTH_RFP **(c)** P_GFP_SpoTH_dCas9 **(d)** P_OL **(e)** P_IFFL_x **(f)** P_Control **(g)** P_GFP_SpoTH_RelA+ **(h)** P_IFFL_RelA_x **(i)** P_BFP

#### Supplementary note 1

The SpoTH gene sequence was constructed based on the BssHII digestion and re-ligation of the spoT gene (pGN19 in [43]), which was shown to only have ppGppasse activity. The digestion and re-ligation of spoT using BssHII introduces a frameshift following the 206 codon and consequently a premature stop codon after the 217 codon. Therefore, the SpoTH sequence only contains the first 217 condons of the product of re-ligating and digesting spoT using BssHII. Finally we modified the initial codon of the endogenous spoT gene from TTG to ATG. The full SpoTH sequence is shown in SI Table 2.

#### Supplementary note 2

##### Summary of simplified SpoTH actuator model

Here, we provide a simplified mathematical model that describes how expressing SpoTH actuates free ribosome concentration. The full model derivation and details can be found in Section: *Derivation of the SpoTH actuator mathematical model*. A key component of the model is the total ribosome concentration equation given by

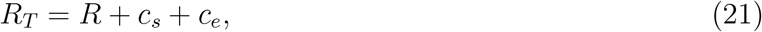

where *R*_*T*_ is the concentration of the total ribosomes in the cell, *R* is the concentration of free ribosomes, and *c*_*s*_ and *c*_*e*_ are the concentrations of the mRNA-ribosome complex corresponding to SpoTH mRNA and the mRNA corresponding to the cell’s endogenous genes, respectively. The concentration of SpoTH is proportional to *c*_*s*_ (3) and thus from hereon, we use varying SpoTH expression and varying *c*_*s*_ interchangeably. Let *c*_*e*_ be a general function of free ribosome concentration, that is, *c*_*e*_ := *c*_*e*_(*R*). We assume that more endogenous mRNA is translated when *R* increases, that is, 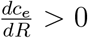. We define

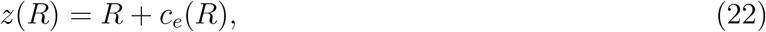

which satisfies *z*(0) = 0 (in the absence of free ribosomes no endegnous mRNA is translated) and is monotonically increasing with *R*. We rewrite (21) using (22), using a model of how *R*_*T*_ depends on *c*_*s*_ through SpoTH catalyzing ppGpp hydrolysis (see Section: *Actuating the ribosomal budget in the cell*), and using overbars to denote concentrations normalized by the total ribosome concentration when there is no ppGpp in the cell, as:

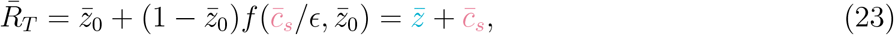

where 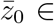 [0, 1] is a proxy of the nominal free ribosome concentration corresponding to no SpoTH expression 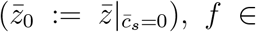 [0, 1) is given by 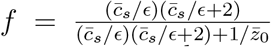 and captures how SpoTH increases the total ribosome concentration, and *ϵ* is a dimensionless parameter that measures how effectively SpoTH catalyzes the hydrolysis of ppGpp and how effectively SpoTH-mRNA is translated into protein. An additional key quantity is

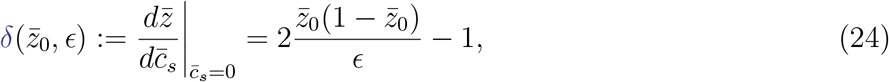

which is the slope of 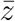 at *c*_*s*_ = 0. The qualitative behavior of (23) is shown in Fig. 18 and it has three qualitatively different responses. When *δ* > 0, we obtain a desired actuator profile where 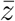 increases initially as 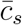 increases, as 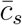 continues to increase the *f* term saturates to unity and the right hand side 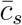 term of (23) dominates and thus the actuator profile peaks and then decreases. As *δ* → ∞, the peak actuation and actuator operational range (as defined in Fig 1-c) both approach the quantity 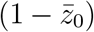. When 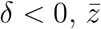 decreases initially as 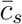 increases and then it can either continue to decrease or it can eventually increase past 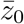, peak, and then decrease again.

###### Remark 1.

From (24), for a fixed *ϵ* such that *ϵ*< 0.5, there exists 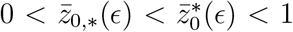 such that for all 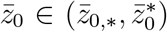 we have that *δ* > 0 and for all 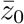 outside this set, *δ* < 0. In (13) we show that 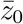 monotonically decreases with basal ppGpp (*θ*_*G*_) and thus for a fixed *ϵ*, there is an open interval of basal ppGpp values that render the desired actuation profile. This implies that too high or too low basal ppGpp can be detrimental in achieving the desired actuator profile.

In Fig. 19, we show the normalized actuation 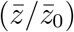 profile for several 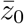 values. We observe that for lower 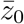 we have more normalized peak actuation. In the inset we show that the normalized peak actuation increases with 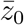 up until 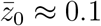. After this critical value, peak actuation decreases as 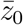 decreases.

**Figure 18:**
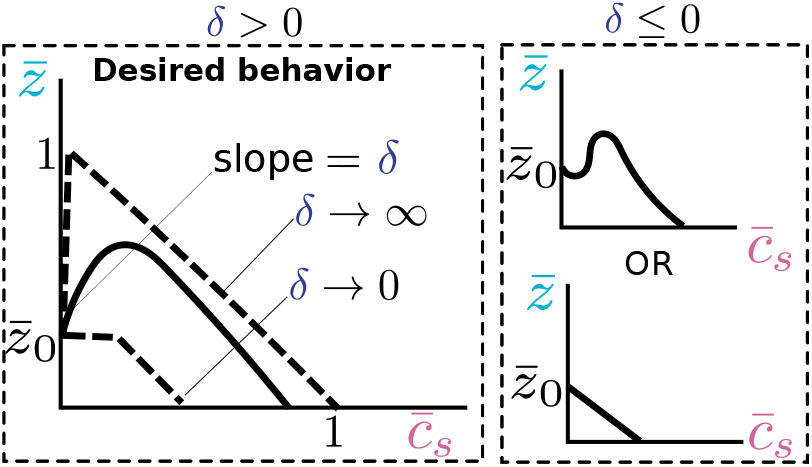
Qualitative behavior of actuator. For *δ* > 0 (24), the actuator profile predicted by (23) has the desired behavior where 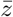 (proxy of free ribosomes) increases as SpoTH is expressed (increasing 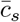), then it peaks and begins to drop. The asymptotic behavior as *δ* → ∞ and *δ* → 0 are depicted by dashed lines. When 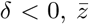 initially decreases as SpoTH is expressed. It can then either continue to decrease or at some point increase, peak, and then decrease again.

**Figure 19:**
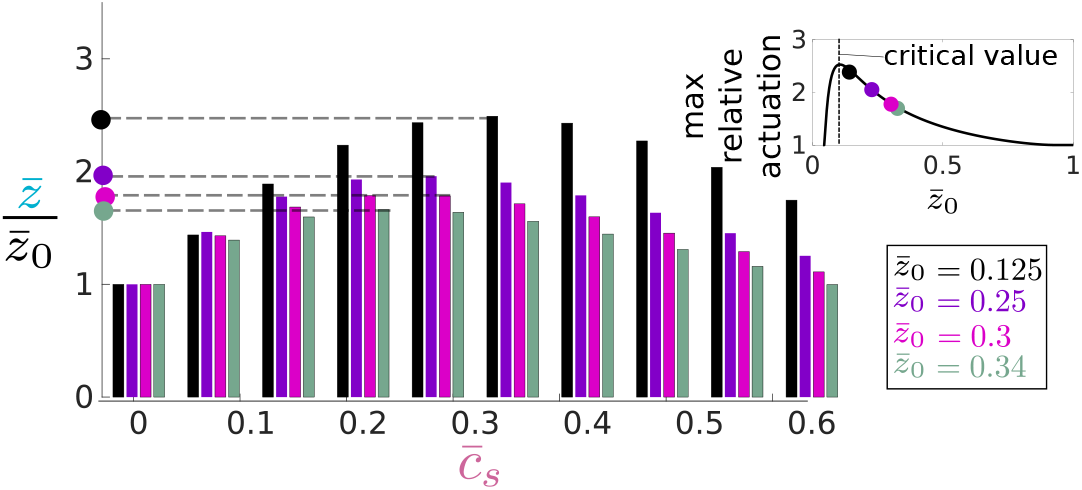
Tradeoff between nominal level and normalized peak actuation. The actuation profile predicted by (23) for several 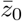. We observe that for lower 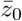, there is higher normalized peak actuation. In the inset we show that the normalized actuation increases as 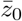 increases up until the critical value of 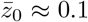. After this critical value normalized peak actuation decreases as 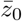 increases. For this simulation we have *ϵ* = 0.13.

##### Simplified model of expressing a heterologous protein

When accounting for the expression of a heterologous protein y, the total ribosome concentration equation (21) is modified to

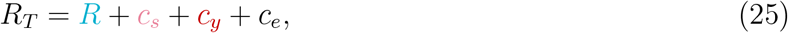

where *c*_*y*_ is the concentration of the mRNA-ribosome complex corresponding to the mRNA of y. The protein concentration of y is proportional to *c*_*y*_. The dimensionless total ribosome equation (23) now reads

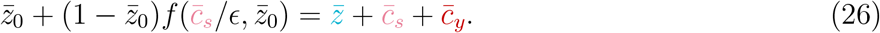

In SI Section: *Derivation of the SpoTH actuator mathematical model*, we show that at steady state, the quantities 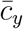 and 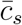 are given by

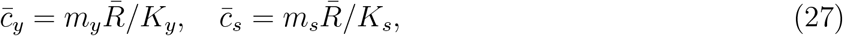

where *m*_*y*_ is the y mRNA concentration, *K*_*y*_ is the dissociation constant of free ribosome with y mRNA, *m*_*s*_ is the SpoTH mRNA concentration, and *K*_*s*_ is the dissociation constant of free ribosome with SpoTH mRNA. Each of *K*_*y*_ and *K*_*s*_ can be tuned by changing the ribosome binding site (RBS) of the corresponding mRNA. From (27), we have that

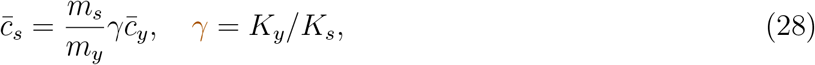

where *γ* is the ratio between the SpoTH RBS strength and the y RBS strength.

##### Feedforward controller

We model SpoTH and y as being transcribed from the same promoter, which implies *m*_*s*_ = *m*_*y*_. We refer to the configuration where y and SpoTH are transcriptionally coupled this way as the closed loop system and it obeys (26) with 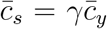 as shown by (28) when *m*_*s*_ = *m*_*y*_. We denote expressing y in the absence of SpoTH as the open loop system and it obeys (26) with 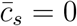 for all values of 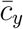. The qualitative behavior of the closed loop system compared to the open loop system is shown in SI Fig. 16. We define the ideal relationship between 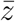 and 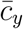 as 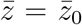for all 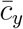, as shown in Fig. 20. The initial slope 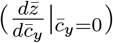 of the closed loop system is given by *γδ* − 1, where *δ* is given by (24). Thus, if *δ* > 0, which, from Fig 18, implies that we are in the parameter regime such that the actuator has a desired profile, then the SpoTH RBS strength (*γ*) can be chosen such that 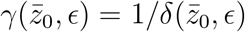 to render an initial flat response of 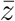 as 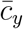 increases. In Fig. 20, we show the closed loop system response (blue lines) for *γ* < 1*/δ, γ* = 1*/δ*, and *γ* > 1*/δ* and the open loop system response (orange). As expected, for *γ* = 1*/δ*, the response of 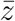 is initially flat as 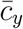 is expressed. The closed loop system achieves higher values of 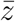 than the open loop system. Furthermore, we observe that the closed loop system achieves higher values of *c*_*y*_ than the open loop system.

**Figure 20:**
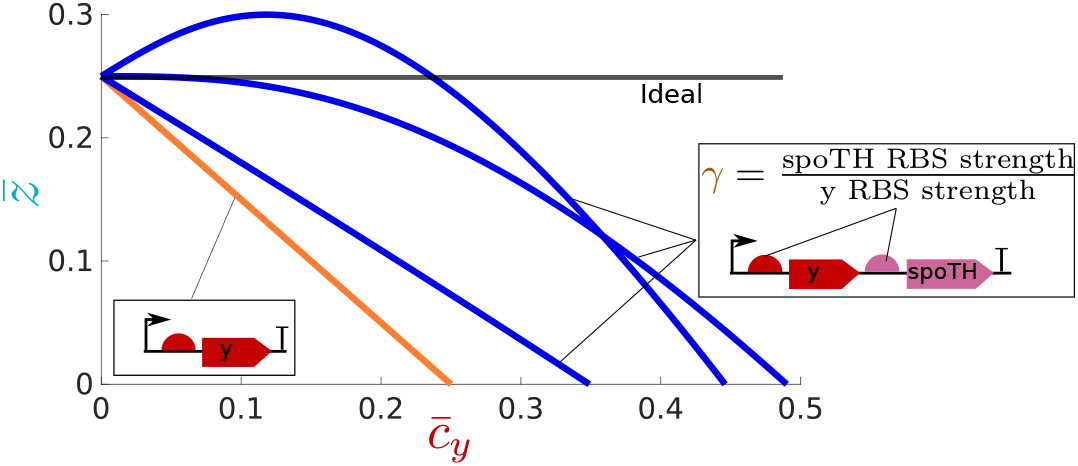
Feedforward controller to compensate for the burden on ribosomes caused by heterologous protein overexpression. Simulation of (26) with 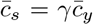. This corresponds to placing y and SpoTH under same promoter (closed loop) depicted in blue. The SpoTH RBS (*γ*) can be tuned to approximate the ideal scenario where 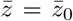for all 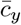. We also show the open loop system (y without SpoTH) depicted in orange as given by (26) with *c*_*s*_ = 0. For this simulation we have 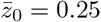 and *ϵ* = 0.13. For the closed loop, we have that *γ* = 0.16, 0.53, 0.9.

#### Supplementary note 3

The model from Supplementary note 2 relates SpoTH expression to free ribosome concentration (or equivalently *z*), here we propose a model to relate SpoTH expression to the cell growth rate (*μ*). A precise model of growth rate as SpoTH is expressed would require a whole-cell model [71]. However, in this work we are interested in the qualitative behavior of growth rate. Thus, we don’t consider an explicit model and rather assume that the growth rate is given by

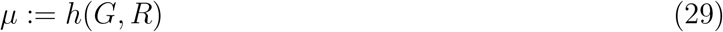

with the properties than 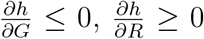, and that *h*(*G*, 0) = 0. The relationship (29) is consistent with the interaction diagram from Fig. 1-a in the main text where ppGpp directly downderegulates growth genes and thus growth rate 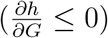 and free ribosome translates mRNA’s responsible for cell growth and thus they upregulate growth rate 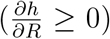. Furthermore, *h*(*G*, 0) = 0 implies that cells cannot grow when there are no free ribosomes present, which is consistent with physical intuition.

##### Growth rate versus the SpoTH gene activation

The change in growth rate as SpoTH is expressed, is given by

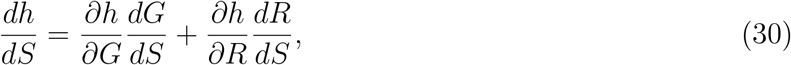

per (6) we have that

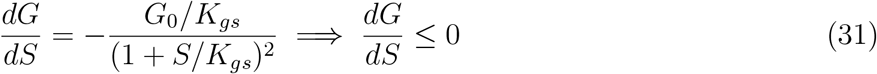

and 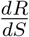 is how free ribosome concentration changes as SpoTH is expressed. From (29) and (31) and our assumptions on *h*, we have that the quantities 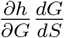 and 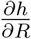 are positive, implying that:

- The mapping between growth rate and SpoTH expression is qualitatively similar to that of Fig. 18 except that the growth rate peak occurs at a higher SpoTH expression level than the peak in free ribosomes. This is consistent with our experimental data where GFP production rate peaked (SI Fig. 7) for lower values of SpoTH (aTc) compared to growth rate (Main text Fig. 2). The data and (30) is also consistent with the fact that growth rate can increase with SpoTH expression while free ribosomes decrease with SpoTH expression. The assumption that *h*(*G*, 0) = 0 implies that the growth rate indeed reaches a maximum when SpoTH is expressed and then approaches zero as the SpoTH mRNA sequesters all the available ribosomes.
- ppGpp levels can be used to tune the mapping between growth rate and SpoTH expression in a similar matter as for free ribosome concentration is tuned (see Remark 1).

##### Growth rate in feedforward controller

In a feedforward configuration we have that the protein level of the GOI (*y*) is proportional to that of SpoTH, that is *y* = *θ*_*y*_*S* (Supplementary note 2), where *θ*_*y*_ is a positive constant. Thus, the change in growth rate as the GOI is expressed in the feedforward configuration is given by

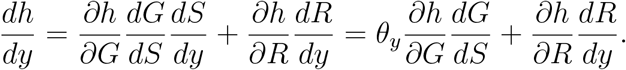

From (31), our assumptions on *h*, and the fact that *θ*_*y*_ > 0, we have that the mapping between growth rate and GOI expression is qualitatively similar to that of Fig. 20 and the SpoTH RBS can be used to make growth rate initially flat as the GOI is expressed. However, the SpoTH RBS that makes growth rate flat 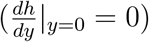 is one where 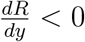, that is, free ribosome decrease with GOI expression. This is consistent with our experimental data where GFP production rate decreases with GOI expression (SI Fig. 9) and growth rate is nearly flat (Main text Fig. 3).

#### Supplementary note 4

Through a simple mathematical model we show that the protein production rate of a constitutive protein is a proxy for free ribosome concentration. This is consistent with [14] where the constitutive expression of a GFP monitor was used as a proxy for free ribosome levels.

We model mRNA (m) binding to free ribosomes (R) to produce the translation initiation complex c, which is then translated to produce the protein P with elongation rate constant *κ*. The mRNA decays with rate constants *δ* and the protein dilutes with rate constant *γ*. The corresponding chemical reactions are:

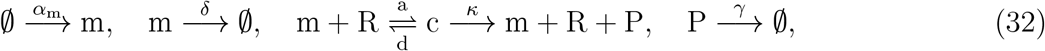

where *α*_*m*_ is the production rate constant of the mRNA, *a* and *d* are the association and dissociation rate constant, respectively, between ribosomes and mRNA. The concentration of each species satisfies:

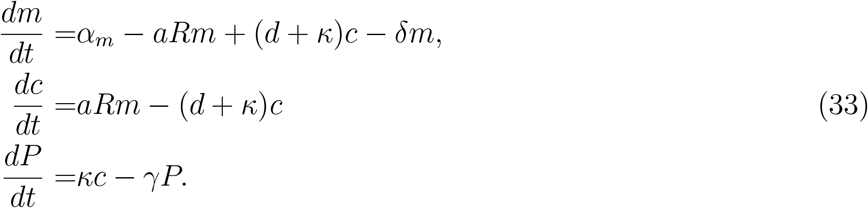

The ribosome-mRNA dynamics can be assumed to be fast relative to *γ* [72] and thus the quasi-steady state [53] of (33) is given by

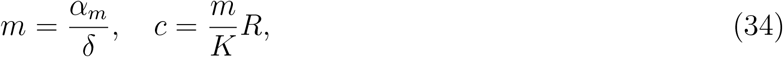

where 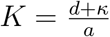. Thus, the reduced protein concentration dynamics are given by

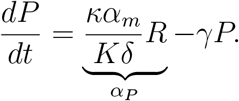

where *α*_*P*_ is the protein production rate. If the protein is constitutively expressed, then *α*_*m*_ is constant and *α*_*P*_ is given by a constant 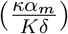 multiplied by *R*, implying that the protein production rate is a proxy for free ribosome concentration.

#### Supplementary note 5

Our modeling framework suggests that we can tune the SpoTH RBS strength in the closed loop genetic circuit (express heterologous protein and SpoTH on the same mRNA) to minimize the sensitivity of free ribosomes on heterologous protein expression (Fig. 20). Therefore, we created a SpoTH RBS library: RBS 1, RBS 2, RBS3, and RBS 4, to test on the closed loop circuit. In this section we characterize the relative strength of the library in the configuration where SpoTH is expressed on the same mRNA as RFP (placed upstream of SpoTH). We show that the strength of the RBS increases in the following order: RBS 1, RBS 2, RBS 3, and RBS 4.

The RBS strength is dependent on the upstream and downstream sequences of the RBS [73, 74], therefore we characterize the SpoTH RBS library with RFP upstream of SpoTH so that the results are applicable to the closed loop controller (Fig. 3-d in the main text). However, we decrease the RBS strength of RFP by several fold (MBP 0.006 in [13]) such that the amount of ribosomes it sequesters are negligible (relative to SpoTH actuation) and thus the change in ribosome concentration when expressing the mRNA with both the weak RFP RBS and SpoTH, is identical to SpoTH in isolation. The construct used to characterize the SpoTH RBS library is shown in Fig.21-a.

Increasing the RBS strength implies that for a fixed amount of SpoTH mRNA, more ribosomes are recruited to translate the mRNA and thus more SpoTH protein is produced. Therefore, less SpoTH mRNA is needed to actuate as the SpoTH RBS strength increases. This implies that when expressing SpoTH using the construct shown in Fig.21-a, less AHL is needed to see an actuation of GFP production rate and growth rate as the SpoTH RBS increases. The GFP production rate and growth rate data are shown in Fig.21-b and Fig.21-c, respectively, when expressing SpoTH using the genetic circuit in Fig.21-a with lactose as the carbon source. We observe that for the list: RBS 1, RBS 2, RBS 3, and RBS 4, that the amount of AHL needed to actuate the GFP production rate and growth rate decreases. Thus, based on our physical intuition, it implies that the RBS strength should have an increasing order of: RBS 1, RBS 2, BS 3, and RBS 4. The same trend is observed in Fig.21-d and Fig.21-e when using glycerol as the carbon source.

Our physical intuition that increasing the SpoTH RBS strength implies that less mRNA is needed to see an actuation on free ribosomes, can be made mathematically precise using the actuator model (12), which relates free ribosome concentration to SpoTH expression. From the fact that

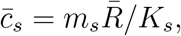

where *m*_*s*_ is the SpoTH mRNA and *K*_*S*_ is inversely proportional to the SpoTH RBS strength, to specify 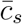 we need to know the value 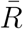. Therefore, we need to specify specify 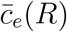. We assume that 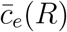 has a form similar to that of 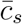, then for *q* different endogenous genes expressing mRNA, 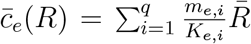, where for gene *i, m*_*e,i*_ is the endogenous mRNA concentration and *K*_*e,i*_ is the effective dissociation constant of endogenous mRNA with ribosomes. In this cases, 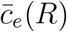 satisfies all of the assumptions stated in Section: *Derivation of the SpoTH actuator mathematical model*. and furthermore, 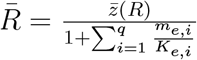. Let 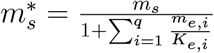 and thus (26) now reads:

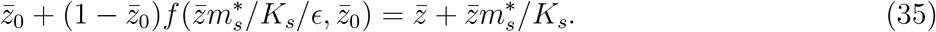

The results from simulating (35) are shown in Fig.22. We observe that increasing the RBS strength (decrease *K*_*s*_) the amount of SpoTH mRNA 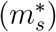 needed to actuate 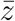 decreases.

**Figure 21:**
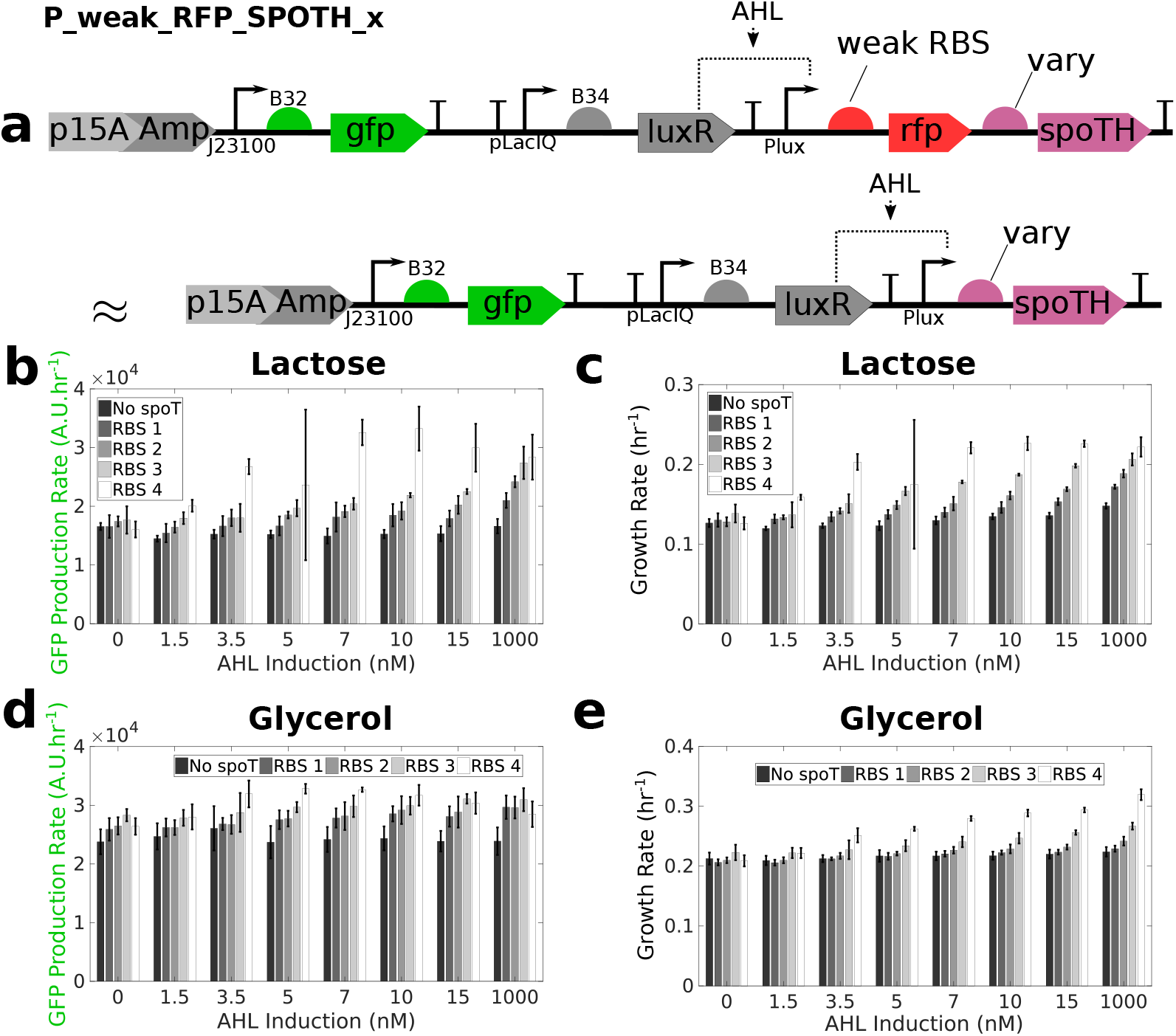
Characterizing the SpoTH RBS library strengths. **(a)** The P_weak_RFP_SPOTH_x genetic construct used to characterize the SpoTH RBS library. This construct is identical to Fig. 3-d (P_IFFL_x) in the main text, but with a very weak RFP RBS strength. The genetic construct P_weak_RFP (identical to P_OL with weak RBS for RFP) corresponds to “No SpoTH” in the legends. Plasmid description, plasmid map, and essential DNA sequences are provided in SI section *Plasmid maps and DNA sequences*. **(b)** For lactose as the carbon source, the GFP production rate as SpoTH is expressed (increase AHL) for the RBS library. **(c)** For lactose as the carbon source, the growth rate as SpoTH is expressed (increase AHL) for the RBS library. **(d)** For glycerol as the carbon source, the GFP production rate as SpoTH is expressed (increase AHL) for the RBS library. **(e)** For glycerol as the carbon source, the growth rate as SpoTH is expressed (increase AHL) for the RBS library. For all data, error bars represent standard deviation from at least four replicates (two biological replicates each with two technical replicates). Data are shown as the mean ± one standard deviation (N=4, two biological replicates each with two technical replicates). All experiments were performed in the CF945 strain. The complete experimental protocol is provided in the Materials and Methods section.

**Figure 22:**
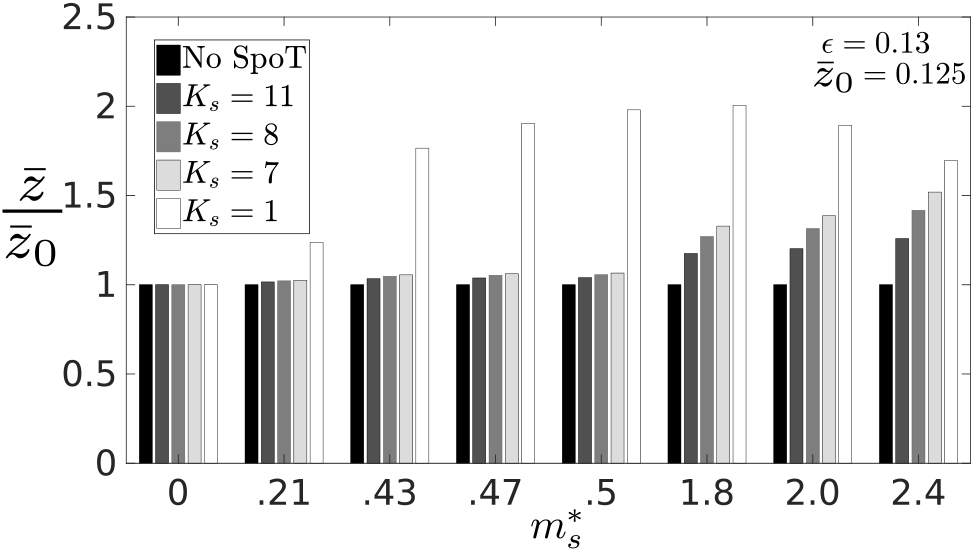
SpoTH expression with several RBS strengths. The normalized measure of free ribosome concentration 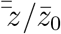 predicted by (35) as the normalized SpoTH mRNA 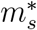 and SpoTH RBS strength (1*/K*_*s*_) are varied. The simulation parameters are *ϵ* = 0.13 and 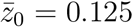. The “No SpoT” bars correspond to 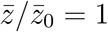 for all 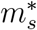 values.

#### Supplementary note 6

For Fig. 5-e in the main text, when the RFP is activated, the growth rate for the OL system in CF946 drops by 40% and that of the associated CL system with SpoTH RBS 2 decreases by 15%. The growth rate for CL RBS 3 corresponding to the set up of Fig. 5-e, monotonically increase by 40% as RFP is activated (SI Fig. 12) thus implying the existence of a CL RBS with strength between RBS 2 and RBS 3 such that the growth rate of the CL systems remains constant as RFP is activated (Fig. 20 and Supplementary Note 3).

#### Supplementary note 7

dCas9 expression is known to be toxic in many bacteria [54, 55]. To this end, we use the SpoTH actuator to reduce growth defects due to overexpressing the dCas9 protein. We express SpoTH using the inducible pTet promoter and dCas9 using the inducible Plux promoter (Fig. 23-a). To estimate the relative production rates of dCas9 between induction values and to assess how much of the burden of expressing dCas9 comes from toxicity rather than ribosome sequestration, we replace dCas9 in Fig. 23-a with RFP (Fig. 23-b). The induction of dCas9 with no SpoTH expression results in a ∼ 40% drop in growth rate (Fig. 23-c). For every dCas9 induction level, there is a SpoTH induction that results in a growth rate that is near the nominal value when no dCas9 nor SpoTH are expressed (colored dots in Fig. 23-c). For AHL = 0.25 nM, growth rate drops by ∼ 8% when expressing dCas9 and without SpoTH, suggesting that toxicity is already present. However, by expressing SpoTH, even for AHL = 2.0 nM, growth rate stays nearly constant (Fig. 23-d). At AHL = 2.0 nM, nearly four times more RFP is produced than at AHL = 0.25 nM (Fig. 23-d). The assumption that RFP production rate is proportional to that of the dCas9, implies that four times more dCas9 is produced at AHL = 2.0 nM than at AHL = 0.25 nM. Thus, we conclude that with the appropriate SpoTH expression, we can produce four times the amount of dCas9 that would otherwise be toxic to the cell, while keeping growth rate constant. Additionally, GFP production rate drops by ∼ 40% when dCas9 is expressed and SpoTH, in principle, can be used to keep it constant (Fig. 23-e,f).

Expressing RFP with the same AHL values as those tested in Fig. 23-c, leads to minimal growth defects but also small drops in GFP production rate when compared to expressing dCas9 (Fig. 23-d,f). Assuming that changes in GFP production rate are a proxy for changes in free ribosome concentration (Supplementary note 4), the incomparable drop in GFP production rate when expressing RFP rather than dCas9 makes it difficult to conclude how much of the burden of expressing dCas9 comes from toxicity rather than ribosome sequestration. To this end, we expressed RFP to a level that would yield a comparable drop in GFP production (more than 40%) as when dCas9 is expressed to the levels in Fig. 23-f, and observed that growth rate only dropped by ∼ 15% (Fig. 24). This indicates that a large portion of the observed growth defects when expressing dCas9 are likely due to direct toxicity as opposed to being due to ribosome sequestration, consistent with published literature [75, 76, 56].

**Figure 23:**
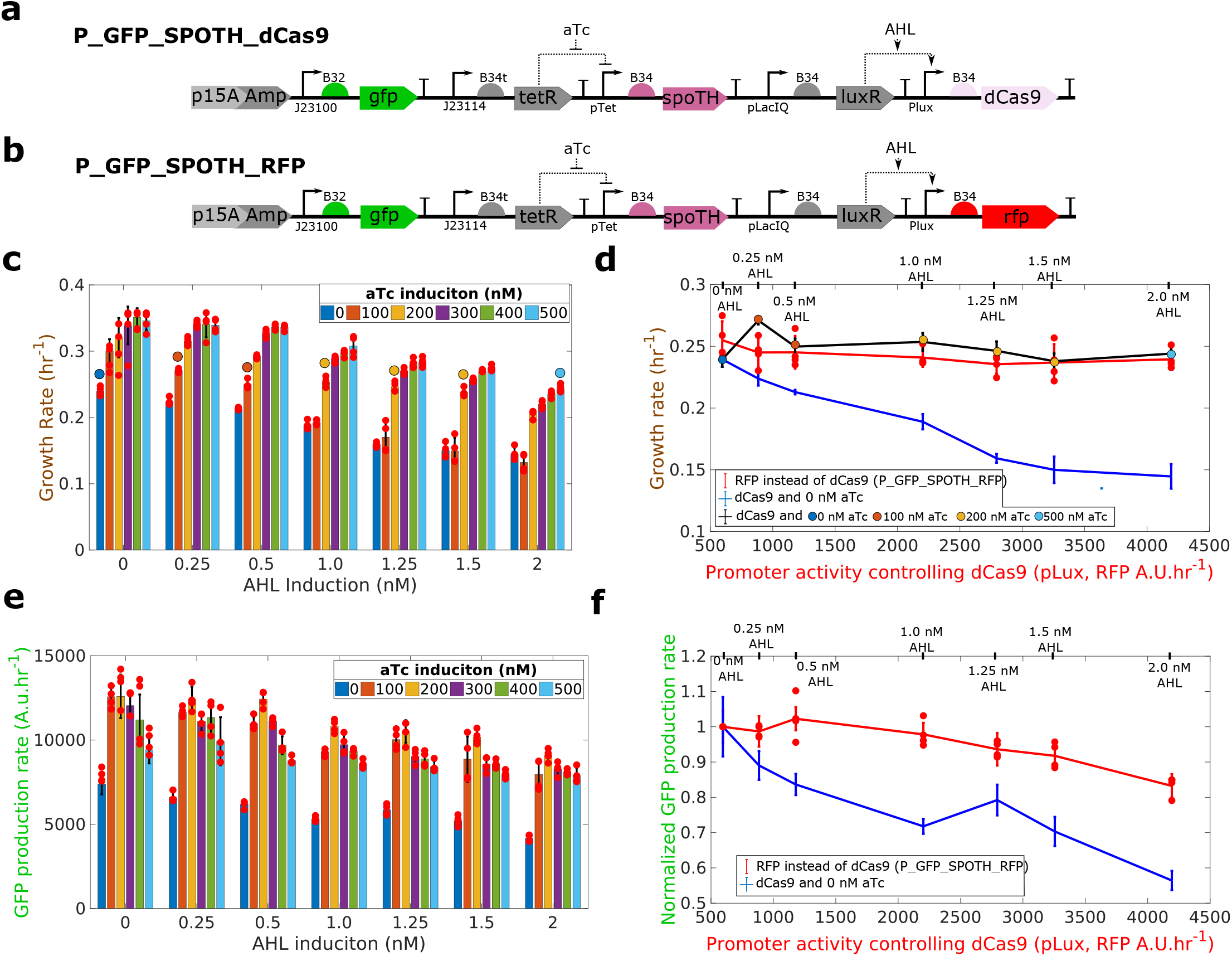
SpoTH actuator rescues growth rate reduction from dCas9 toxicity. **(a)** P_GFP_SpoTH_dCas9 plasmid used to express SpoTH using the inducible pTet promoter and dCas9 using the inducible Plux promoter. **(b)** P_GFP_SpoTH_RFP plasmid used to express SpoTH using the inducible pTet promoter and RFP using the inducible Plux promoter. **(c)** Cell growth rate as dCas9 production rate is increased via AHL, for different levels of SpoTH induction. For each AHL we mark by a circle the aTc level that results in a growth rate closest to the nominal growth rate when AHL = 0 nM and aTc = 0 nM. **(d)** Cell growth rate versus pLux promoter activity as measured in units of RFP production rate for the same AHL induction levels as in (c) when pLux is transcribing RFP (red line, P_GFP_SpoTH_RFP with no SpoTH induction), dCas9 and with no SpoTH induction (blue line), and dCas9 with SpoTH induction corresponding to the colored circles in (c) (black line). **(e)** GFP production rate as dCas9 production rate is increased via AHL, for different levels of SpoTH induction. **(f)** GFP production rate versus pLux promoter activity as measured in units of RFP production rate for the same AHL induction levels as in (e) when pLux is transcribing RFP (red line, P_GFP_SpoTH_RFP with no SpoTH induction) and dCas9 and with no SpoTH induction (blue line). Data are shown as the mean ± one standard deviation (N=4, two biological replicates each with two technical replicates). Individual experimental values are presented as a red dots. All experiments were performed in the CF945 strain in media with glycerol as the sole carbon source. Plasmid description, plasmid map, and essential DNA sequences are provided in SI section *Plasmid maps and DNA sequences*. The complete experimental protocol is provided in the Materials and Methods section.

**Figure 24:**
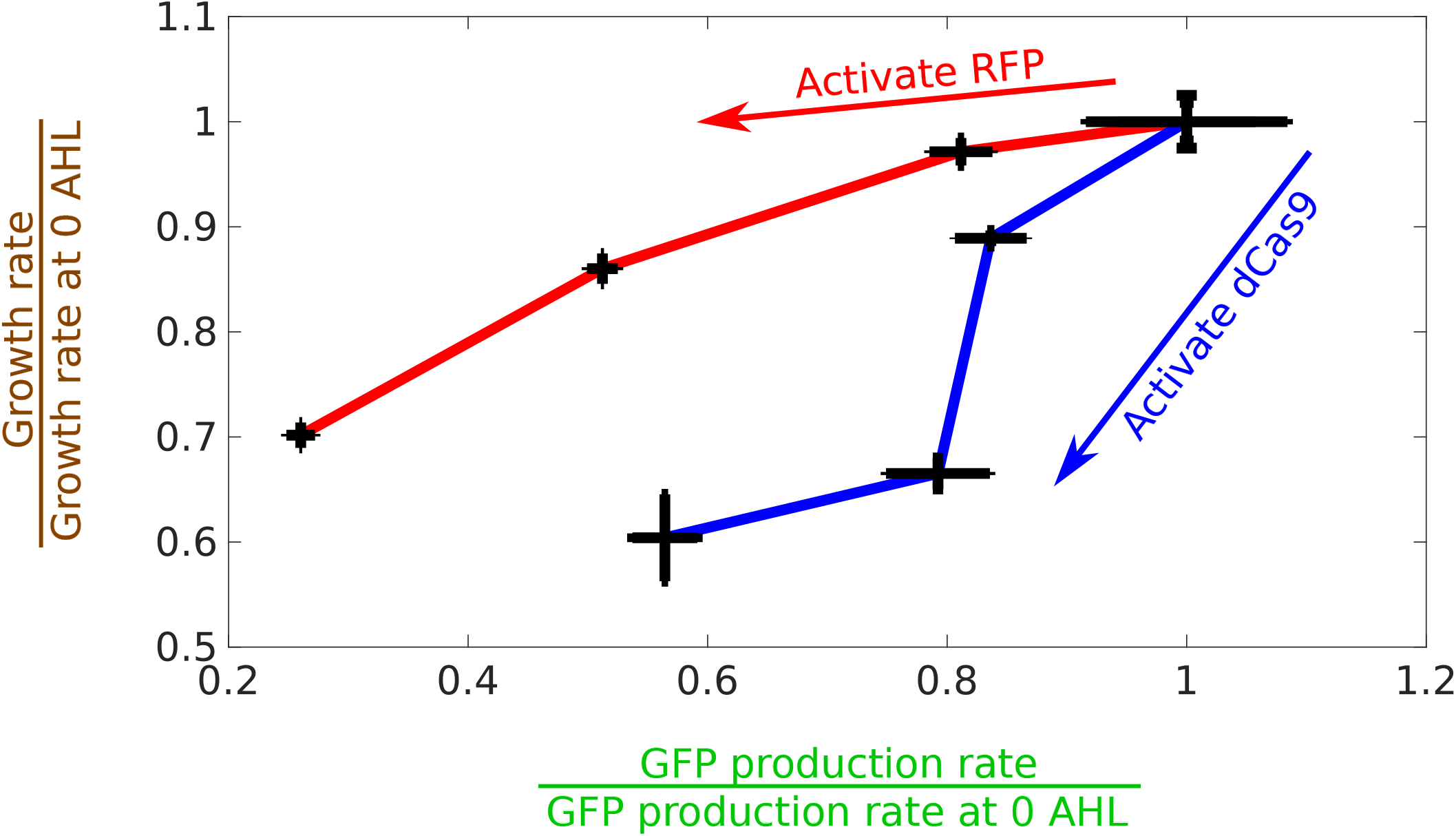
dCas9 expression places more growth rate burden on the cell than RFP expression due to toxic effects. dCas9 and RFP are expressed using the P_GFP_SpoTH_dCas9 (blue line) and P_GFP_SpoTH_RFP (red line) plasmids, respectively (SI Fig. 23). The GFP production rate normalized by GFP production rate when there is no AHL induction versus the growth rate normalized by growth rate when there is no AHL induction. The inductions for P_GFP_SpoTH_dCas9 are 0 nM, 0.5 nM, 1.25 nM, and 2 nM AHL and for P_GFP_SpoTH_RFP are 0 nM, 3 nM, 7 nM, 15 nM AHL. Data are shown as the mean ± one standard deviation (N=4, two biological replicates each with two technical replicates). All experiments were performed in the CF945 strain in media with glycerol as the sole carbon source. The complete experimental protocol is provided in the Materials and Methods section.

## References

[1] J. Claesen and M. A. Fischbach, “Synthetic Microbes As Drug Delivery Systems,” ACS Synth. Biol., vol. 4, pp. 358–364, apr 2015.

[2] R. M. Shia, L. K. McIntire, J. A. Hagen, C. D. Goodyear, L. N. Dykstra, and A. R. Myers, “Biomarker and Biometric Indices of Physical Exhaustion in the Firefighting Community,” Procedia Manuf., vol. 3, pp. 5081–5087, 2015.

[3] J. Dou and M. R. Bennett, “Synthetic Biology and the Gut Microbiome,” Biotechnol. J., vol. 13, p. 1700159, may 2018.

[4] A. Cubillos-Ruiz, M. A. Alcantar, N. M. Donghia, P. Cárdenas, J. Avila-Pacheco, and J. J. Collins, “An engineered live biotherapeutic for the prevention of antibiotic-induced dysbiosis,” Nat. Biomed. Eng., apr 2022.

[5] P. Dvorák, P. I. Nikel, J. Damborský, and V. de Lorenzo, “Bioremediation 3. 0 : Engineering pollutant-removing bacteria in the times of systemic biology,” Biotechnol. Adv., vol. 35, pp. 845– 866, nov 2017.

[6] W. Thavarajah, M. S. Verosloff, J. K. Jung, K. K. Alam, J. D. Miller, M. C. Jewett, S. L. Young, and J. B. Lucks, “A primer on emerging field-deployable synthetic biology tools for global water quality monitoring,” npj Clean Water, vol. 3, p. 18, dec 2020.

[7] B. Saltepe, L. Wang, and B. Wang, “Synthetic biology enables field-deployable biosensors for water contaminants,” TrAC Trends Anal. Chem., vol. 146, p. 116507, jan 2022.

[8] I. Del Valle, E. M. Fulk, P. Kalvapalle, J. J. Silberg, C. A. Masiello, and L. B. Stadler, “Translating New Synthetic Biology Advances for Biosensing Into the Earth and Environmental Sciences,” Front. Microbiol., vol. 11, feb 2021.

[9] C. A. Voigt, “Synthetic biology 2020–2030: six commercially-available products that are changing our world,” Nat. Commun., vol. 11, p. 6379, ec 2020.

[10] H. L. Pham, H. Ling, and M. W. Chang, “Design and fabrication of field-deployable microbial biosensing devices,” Curr. Opin. Biotechnol., vol. 76, p. 102731, aug 2022.

[11] I. Shachrai, A. Zaslaver, U. Alon, and E. Dekel, “Cost of Unneeded Proteins in E. coli Is Reduced after Several Generations in Exponential Growth,” Mol. Cell, vol. 38, no. 5, pp. 758–767, 2010.

[12] M. S. Bienick, K. W. Young, J. R. Klesmith, E. E. Detwiler, K. J. Tomek, and T. A. Whitehead, “The interrelationship between promoter strength, gene expression, and growth rate,” PLoS One, vol. 9, no. 10, 2014.

[13] A. Gyorgy, J. I. Jiménez, J. Yazbek, H. H. Huang, H. Chung, R. Weiss, and D. Del Vecchio, “Isocost Lines Describe the Cellular Economy of Genetic Circuits,” Biophys. J., vol. 109, no. 3, pp. 639–646, 2015.

[14] F. Ceroni, R. Algar, G.-B. Stan, and T. Ellis, “Quantifying cellular capacity identifies gene expression designs with reduced burden,” Nat. Methods, vol. 12, no. 5, pp. 415–418, 2015.

[15] Y. Qian, H. H. Huang, J. I. Jiménez, and D. Del Vecchio, “Resource Competition Shapes the Response of Genetic Circuits,” ACS Synth. Biol., vol. 6, no. 7, pp. 1263–1272, 2017.

[16] R. Zhang, J. Li, J. Melendez-Alvarez, X. Chen, P. Sochor, H. Goetz, Q. Zhang, T. Ding, X. Wang, and X.-J. Tian, “Topology-dependent interference of synthetic gene circuit function by growth feedback,” Nat. Chem. Biol., vol. 16, pp. 695–701, jun 2020.

[17] R. Zhang, H. Goetz, J. Melendez-Alvarez, J. Li, T. Ding, X. Wang, and X. J. Tian, “Winnertakes-all resource competition redirects cascading cell fate transitions,” Nat. Commun., vol. 12, no. 1, pp. 1–9, 2021.

[18] S. R. Scott, M. O. Din, P. Bittihn, L. Xiong, L. S. Tsimring, and J. Hasty, “A stabilized microbial ecosystem of self-limiting bacteria using synthetic quorum-regulated lysis,” Nat. Microbiol., vol. 2, no. June, pp. 1–9, 2017.

[19] A. Miano, M. J. Liao, and J. Hasty, “Inducible cell-to-cell signaling for tunable dynamics in microbial communities,” Nat. Commun., vol. 11, no. 1, 2020.

[20] A. J. Fedorec, B. D. Karkaria, M. Sulu, and C. P. Barnes, “Single strain control of microbial consortia,” Nat. Commun., vol. 12, no. 1, pp. 1–12, 2021.

[21] T. Shopera, L. He, T. Oyetunde, Y. J. Tang, and T. S. Moon, “Decoupling Resource-Coupled Gene Expression in Living Cells,” ACS Synth. Biol., vol. 6, no. 8, pp. 1596–1604, 2017.

[22] H. H. Huang, Y. Qian, and D. Del Vecchio, “A quasi-integral controller for adaptation of genetic modules to variable ribosome demand,” Nat. Commun., vol. 9, no. 1, pp. 1–12, 2018.

[23] R. D. Jones, Y. Qian, V. Siciliano, B. DiAndreth, J. Huh, R. Weiss, and D. Del Vecchio, “An endoribonuclease-based feedforward controller for decoupling resource-limited genetic modules in mammalian cells,” Nat. Commun., vol. 11, no. 1, pp. 1–16, 2020.

[24] R. D. Jones, Y. Qian, K. Ilia, B. Wang, M. T. Laub, D. Del Vecchio, and R. Weiss, “Robust and tunable signal processing in mammalian cells via engineered covalent modification cycles,” Nat. Commun., vol. 13, p. 1720, ec 2022.

[25] T. Frei, F. Cella, F. Tedeschi, J. Gutiérrez, G. B. Stan, M. Khammash, and V. Siciliano, “Charac-terization and mitigation of gene expression burden in mammalian cells,” Nat. Commun., vol. 11, no. 1, pp. 1–14, 2020.

[26] A. P. Darlington, J. Kim, J. I. Jiménez, and D. G. Bates, “Dynamic allocation of orthogonal ribosomes facilitates uncoupling of co-expressed genes,” Nat. Commun., vol. 9, no. 1, 2018.

[27] F. Ceroni, A. Boo, S. Furini, T. E. Gorochowski, O. Borkowski, Y. N. Ladak, A. R. Awan, C. Gilbert, G. B. Stan, and T. Ellis, “Burden-driven feedback control of gene expression,” Nat. Methods, vol. 15, no. 5, pp. 387–393, 2018.

[28] J. A. N. Brophy and C. A. Voigt, “Principles of genetic circuit design,” Nat. Methods, vol. 11, no. 5, pp. 508–520, 2014.

[29] T. E. Gorochowski, A. Espah Borujeni, Y. Park, A. A. Nielsen, J. Zhang, B. S. Der, D. B. Gordon, and C. A. Voigt, “Genetic circuit characterization and debugging using RNA -seq, ” Mol. Syst. Biol., vol. 13, no. 11, p. 952, 2017.

[30] B. J. Paul, W. Ross, T. Gaal, and R. L. Gourse, “rRNA transcription in Escherichia coli,” Annu. Rev. Genet., vol. 38, pp. 749–770, 2004.

[31] K. Potrykus, H. Murphy, N. Philippe, and M. Cashel, “ppGpp is the major source of growth rate control in E. coli,” Environ. Microbiol., vol. 13, no. 3, pp. 563–575, 2011.

[32] N. C. Imholz, M. J. Noga, N. J. van den Broek, and G. Bokinsky, “Calibrating the Bacterial Growth Rate Speedometer: A Re-evaluation of the Relationship Between Basal ppGpp, Growth, and RNA Synthesis in Escherichia coli,” Front. Microbiol., vol. 11, no. September, pp. 1–9, 2020.

[33] G. Schreiber, S. Metzger, E. Aizenman, S. Roza, M. Cashel, and G. Glaser, “Overexpression of the relA gene in Escherichia coli,” J. Biol. Chem., vol. 266, pp. 3760–3767, feb 1991.

[34] A. Svitil, M. Cashel, and J. Zyskind, “Guanosine tetraphosphate inhibits protein synthesis in vivo. A possible protective mechanism for starvation stress in Escherichia coli.,” J. Biol. Chem., vol. 268, pp. 2307–2311, feb 1993.

[35] E. Sarubbi, K. E. Rudd, and M. Cashel, “Basal ppGpp level adjustment shown by new spoT mutants affect steady state growth rates and rrnA ribosomal promoter regulation in Escherichia coli,” MGG Mol. Gen. Genet., vol. 213, no. 2-3, pp. 214–222, 1988.

[36] M. Zacharias, H. U. Goringer, and R. Wagner, “Influence of the GCGC discriminator motif introduced into the ribosomal RNA P2- and tac promoter on growth-rate control and stringent sensitivity,” EMBO J., vol. 8, no. 11, pp. 3357–3363, 1989.

[37] P. P. Dennis and H. Bremer, “Modulation of Chemical Composition and Other Parameters of the Cell at Different Exponential Growth Rates,” EcoSal Plus, vol. 3, no. 1, 2008.

[38] X. Dai and M. Zhu, “Coupling of Ribosome Synthesis and Translational Capacity with Cell Growth,” Trends Biochem. Sci., vol. 45, pp. 681–692, aug 2020.

[39] G. C. Atkinson, T. Tenson, and V. Hauryliuk, “The RelA/SpoT Homolog (RSH) superfamily: Distribution and functional evolution of ppgpp synthetases and hydrolases across the tree of life,” PLoS One, vol. 6, no. 8, 2011.

[40] V. Hauryliuk, G. C. Atkinson, K. S. Murakami, T. Tenson, and K. Gerdes, “Recent functional insights into the role of (p)ppGpp in bacterial physiology,” Nat. Rev. Microbiol., vol. 13, no. 5, pp. 298–309, 2015.

[41] L. Fernández-Coll and M. Cashel, “Possible Roles for Basal Levels of (p)ppGpp: Growth Efficiency Vs. Surviving Stress,” Front. Microbiol., vol. 11, no. October, 2020.

[42] K. D. Murray and H. Bremer, “Control of spoT-dependent ppGpp Synthesis and Degradation in Escherichia coli,” J. Mol. Biol, vol. 259, pp. 41–57, 1996.

[43] D. R. Gentry and M. Cashel, “Mutational analysis of the Escherichia coli spoT gene identifies distinct but overlapping regions involved in ppGpp synthesis and degradation,” Mol. Microbiol., vol. 19, no. 6, pp. 1373–1384, 1996.

[44] V. J. Hernandez and H. Bremer, “Guanosine tetraphosphate (ppGpp) dependence of the growth rate control of rrnB P1 promoter activity in Escherichia coli,” J. Biol. Chem., vol. 265, no. 20, pp. 11605–11614, 1990.

[45] H. Xiao, M. Kalman, K. Ikehara, S. Zemel, G. Glaser, and M. Cashel, “Residual guanosine 3’,5’-bispyrophosphate synthetic activity of relA null mutants can be eliminated by spoT null mutations,” J. Biol. Chem., vol. 266, no. 9, pp. 5980–5990, 1991.

[46] P. P. Dennis and H. Bremer, “Modulation of Chemical Composition and Other Parameters of the Cell at Different Exponential Growth Rates,” EcoSal Plus, vol. 3, no. 1, 2008.

[47] V. J. Hernandez and H. Bremer, “Characterization of RNA and DNA synthesis in Escherichia coli strains devoid of ppGpp,” J. Biol. Chem., vol. 268, no. 15, pp. 10851–10862, 1993.

[48] M. Zhu, H. Mu, M. Jia, L. Deng, and X. Dai, “Control of ribosome synthesis in bacteria: the important role of rRNA chain elongation rate,” Sci. China Life Sci., vol. 3, pp. 169–84, aug 2020.

[49] M. C. Chan, S. Karasawa, H. Mizuno, I. Bosanac, D. Ho, G. G. Privé, A. Miyawaki, and M. Ikura, “Structural characterization of a blue chromoprotein and its yellow mutant from the sea anemone Cnidopus japonicus,” J. Biol. Chem., vol. 281, no. 49, pp. 37813–37819, 2006.

[50] R. Tsoi, F. Wu, C. Zhang, S. Bewick, D. Karig, and L. You, “Metabolic division of labor in microbial systems,” Proc. Natl. Acad. Sci. U. S. A., vol. 115, no. 10, pp. 2526–2531, 2018.

[51] C. V. Dinh, X. Chen, and K. L. Prather, “Development of a Quorum-Sensing Based Circuit for Control of Coculture Population Composition in a Naringenin Production System,” ACS Synth. Biol., vol. 9, no. 3, pp. 590–597, 2020.

[52] M. Zhu, Y. Pan, and X. Dai, “(p)ppGpp: the magic governor of bacterial growth economy,” Curr. Genet., vol. 65, no. 5, pp. 1121–1125, 2019.

[53] D. Del Vecchio and R. M. Murray, Biomolecular Feedback Systems. 2014.

[54] J. M. Rock, F. F. Hopkins, A. Chavez, M. Diallo, M. R. Chase, E. R. Gerrick, J. R. Pritchard, G. M. Church, E. J. Rubin, C. M. Sassetti, D. Schnappinger, and S. M. Fortune, “Programmable transcriptional repression in mycobacteria using an orthogonal CRISPR interference platform,” Nat. Microbiol., vol. 2, no. February, pp. 1–9, 2017.

[55] Y. J. Lee, A. Hoynes-O’Connor, M. C. Leong, and T. S. Moon, “Programmable control of bacterial gene expression with the combined CRISPR and antisense RNA system,” Nucleic Acids Res., vol. 44, no. 5, pp. 2462–2473, 2016.

[56] S. Zhang and C. A. Voigt, “Engineered dCas9 with reduced toxicity in bacteria: Implications for genetic circuit design,” Nucleic Acids Res., vol. 46, no. 20, pp. 11115–11125, 2018.

[57] J. Fontana, C. Dong, J. Y. Ham, J. G. Zalatan, and J. M. Carothers, “Regulated Expression of sgRNAs Tunes CRISPRi in E. coli,” Biotechnol. J., vol. 13, p. 1800069, sep 2018.

[58] H. H. Huang, M. Bellato, Y. Qian, P. Cárdenas, L. Pasotti, P. Magni, and D. Del Vecchio, “dCas9 regulator to neutralize competition in CRISPRi circuits,” Nat. Commun., vol. 12, no. 1, pp. 1–7, 2021.

[59] D. Sun, G. Lee, J. H. Lee, H. Y. Kim, H. W. Rhee, S. Y. Park, K. J. Kim, Y. Kim, B. Y. Kim, J. I. Hong, C. Park, H. E. Choy, J. H. Kim, Y. H. Jeon, and J. Chung, “A metazoan ortholog of SpoT hydrolyzes ppGpp and functions in starvation responses,” Nat. Struct. Mol. Biol., vol. 17, no. 10, pp. 1188–1194, 2010.

[60] M. Zhu and X. Dai, “Growth suppression by altered (p)ppGpp levels results from non-optimal resource allocation in Escherichia coli,” Nucleic Acids Res., vol. 47, no. 9, pp. 4684–4693, 2019.

[61] R. Harinarayanan, H. Murphy, and M. Cashel, “Synthetic growth phenotypes of Escherichia coli lacking ppGpp and transketolase A (tktA) are due to ppGpp-mediated transcriptional regulation of tktB,” Mol. Microbiol., vol. 69, pp. 882–894, aug 2008.

[62] F. Ceroni, R. Algar, G.-B. Stan, and T. Ellis, “Quantifying cellular capacity identifies gene expression designs with reduced burden,” Nat. Methods, vol. 12, no. 5, pp. 415–418, 2015.

[63] D. G. Gibson, L. Young, R. Y. Chuang, J. C. Venter, C. A. Hutchison, and H. O. Smith, “Enzymatic assembly of DNA molecules up to several hundred kilobases,” Nat. Methods, vol. 6, no. 5, pp. 343–345, 2009.

[64] D. Del Vecchio, R. M. Murray, and D. Vecchio, Biomolecular Feedback Systems. 2010.

[65] K. Potrykus and M. Cashel, “Growth at best and worst of times,” Nat. Microbiol., vol. 3, no. 8, pp. 862–863, 2018.

[66] A. G. Marr, “Growth Rate of Escherichia coli,” Microbiol. Mol. Biol. Rev., vol. 55, no. 2, pp. 316– 333, 1991.

[67] H. D. Murray, J. A. Appleman, and R. L. Gourse, “Regulation of the Escherichia coli rrnB P2 promoter,” J. Bacteriol., vol. 185, no. 1, pp. 28–34, 2003.

[68] M. M. Barker, T. Gaal, C. A. Josaitis, and R. L. Gourse, “Mechanism of regulation of transcription initiation by ppGpp. I. Effects of ppGpp on transcription initiation in vivo and in vitro,” J. Mol. Biol., vol. 305, no. 4, pp. 673–688, 2001.

[69] P. P. Dennis, M. Ehrenberg, and H. Bremer, “Control of rRNA synthesis in Escherichia coli: a systems biology approach.,” Microbiol Mol Biol Rev, vol. 68, no. 4, pp. 639–668, 2004.

[70] C. D. McBride and D. Del Vecchio, “Predicting Composition of Genetic Circuits with Resource Competition: Demand and Sensitivity,” ACS Synth. Biol., vol. 10, pp. 3330–3342, ec 2021.

[71] J. Carrera and M. W. Covert, “Why Build Whole-Cell Models?,” Trends Cell Biol., vol. 25, pp. 719–722, ec 2015.

[72] Y. Qian and D. Del Vecchio, “Mitigation of ribosome competition through distributed sRNA feedback,” 2016 IEEE 55th Conf. Decis. Control. CDC 2016, pp. 758–763, 2016.

[73] H. M. Salis, E. A. Mirsky, and C. A. Voigt, “Automated design of synthetic ribosome binding sites to control protein expression,” Nat. Biotechnol., 2009.

[74] A. Espah Borujeni, A. S. Channarasappa, and H. M. Salis, “Translation rate is controlled by coupled trade-offs between site accessibility, selective RNA unfolding and sliding at upstream standby sites,” Nucleic Acids Res., vol. 42, no. 4, pp. 2646–2659, 2014.

[75] P. D. Hsu, D. A. Scott, J. A. Weinstein, F. A. Ran, S. Konermann, V. Agarwala, Y. Li, E. J. Fine, X. Wu, O. Shalem, T. J. Cradick, L. A. Marraffini, G. Bao, and F. Zhang, “DNA targeting specificity of RNA-guided Cas9 nucleases,” Nat. Biotechnol., vol. 31, no. 9, pp. 827–832, 2013.

[76] S. H. Sternberg, S. Redding, M. Jinek, E. C. Greene, and J. A. Doudna, “DNA interrogation by the CRISPR RNA-guided endonuclease Cas9,” Nature, vol. 507, no. 7490, pp. 62–67, 2014.

